# Fast And Accurate Population Level Transcranial Magnetic Stimulation via Low-Rank Probabilistic Matrix Decomposition (PMD)

**DOI:** 10.1101/2023.02.08.527758

**Authors:** Nahian I. Hasan, Dezhi Wang, Luis J. Gomez

## Abstract

Transcranial magnetic stimulation (TMS) is used to study brain function and treat mental health disorders. During TMS, a coil placed on the scalp induces an E-field in the brain that modulates its activity. TMS is known to stimulate regions that are exposed to a large E-field. Clinical TMS protocols prescribe a coil placement based on scalp landmarks. There are inter-individual variations in brain anatomy that result in variations in the TMS-induced E-field at the targeted region and its outcome. These variations across individuals could in principle be minimized by developing a large database of head subjects and determining scalp landmarks that maximize E-field at the targeted brain region while minimizing its variation using computational methods. However, this approach requires repeated execution of a computational method to determine the E-field induced in the brain for a large number of subjects and coil placements. We developed a probabilistic matrix decomposition-based approach for rapidly evaluating the E-field induced during TMS for a large number of coil placements. Our approach can determine the E-field induced in over 1 Million coil placements in 9.5 hours, in contrast, to over 5 years using a bruteforce approach. After the initial set-up stage, the E-field can be predicted over the whole brain within 2-3 milliseconds and to 2% accuracy. We tested our approach in over 200 subjects and achieved an error of < 2% in most and < 3.5% in all subjects. We will present several examples of bench-marking analysis for our tool in terms of accuracy and speed across and its applicability for population level optimization of coil placement.

**Graphical Abstract:** 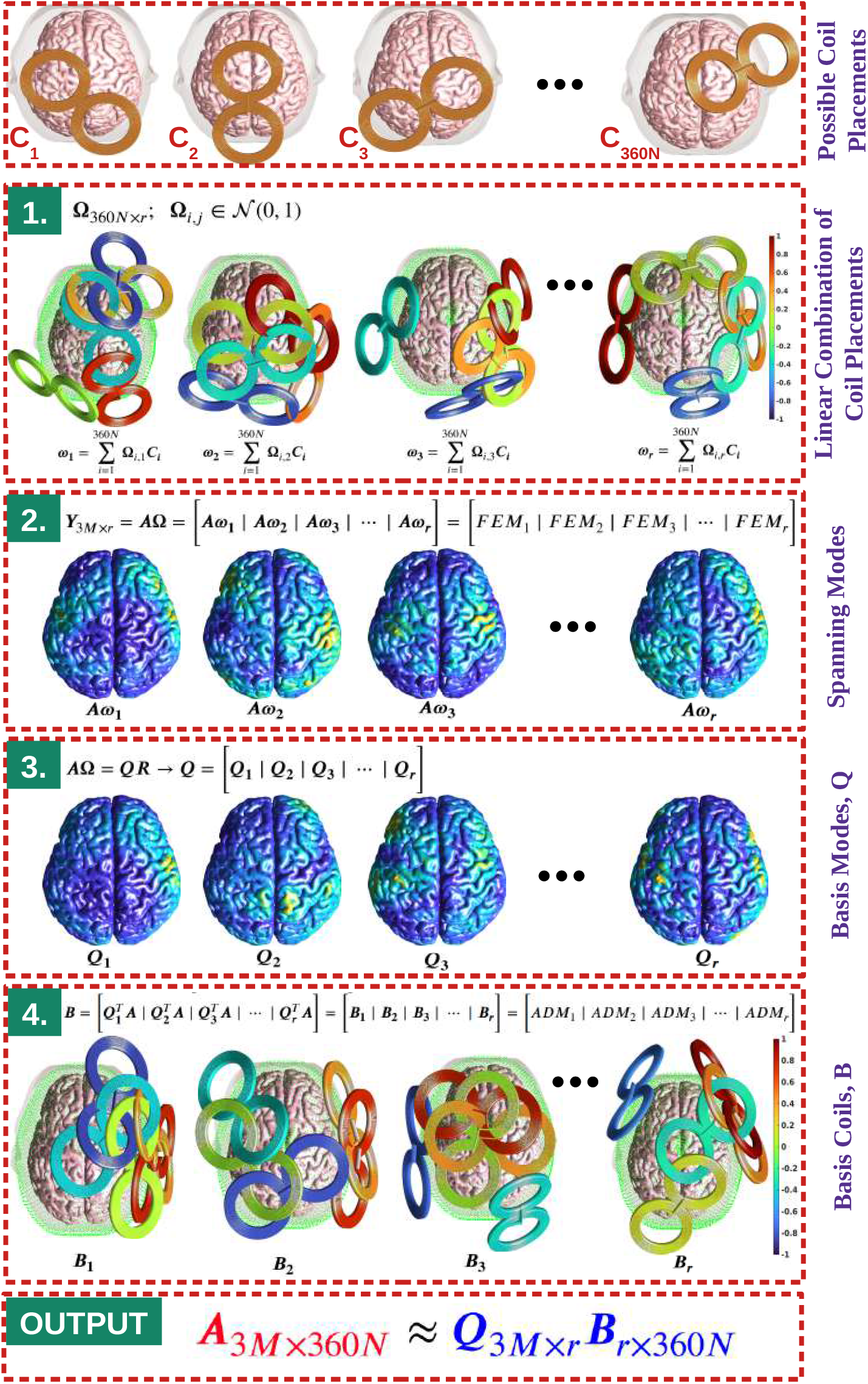

**Highlights:**

- A method for practical E-field informed population-level TMS coil placement strategies is developed.
- This algorithm enables the determination of E-field informed optimal coil placement in seconds enabling it’s use for close-loop and on-the-fly reconfiguration of TMS.
- After the initial set-up stage of less than 10 hours, the E-field can be predicted for any coil placement across the whole brain within in 2-3 milliseconds.

## 1. Introduction

Transcranial Magnetic Stimulation (TMS) is a non-invasive brain stimulation modality where a coil is used to induce an E-field in the brain and modulate its activity [1,2]. The intensity of the induced E-field in the brain has been shown to be an effective predictor of cortical regions that are directly stimulated during TMS [3,4]. To ensure that the targeted cortical site is stimulated, it is important to place the coil where it induces a maximum E-field in it. This work proposes a fast computational technique based on a probabilistic low-rank matrix approximation algorithm [5] that enables finding the E-field for an arbitrary coil placement in near real-time scenarios. Furthermore, we use our solver along with fMRI and EEG pipelines that co-register subjects to develop a population of head models and determine population-level E-field-informed optimum coil placements.

If a head and coil model is available, the E-field induced during TMS can be determined by using the finite difference method (FDM) [6, 7], finite element method (FEM) [8, 9], admittance method [10], or boundary element method (BEM) [11, 12]. These methods predict the E-field distribution inside the head for a given coil placement. However, in many clinical applications of TMS, a head model is not readily available and scalp landmarks are used to determine the coil position in lieu of E-field dose information [13, 14, 15, 16]. These scalp landmarks are oftentimes not aligned with targeted cortical regions [17, 18]. In principle, these landmarks could be determined by running simulations on a population of head models and determining an E-field-informed optimum scalp landmark. This would require a population of head models with co-registered brains and scalps. Furthermore, it would require repeated execution for a large number of coil placements and head models of computational E-field dosimetry software. This is currently computationally intractable using standard FEM, BEM, and FDM techniques. For example, if we choose 2500 possible coil locations on the scalp each associated with 360 coil orientations, assuming 170 seconds per simulation (based on our in-house FEM solver to give < 2% error), this would take approximately 5 years per subject to predict the E-field over the whole brain for all coil placements. This could be lowered by improving the implementation or by lowering the accuracy of the solver. However, even with code optimizations, determining the E-field in a standard resolution of the head model in SimNIBS takes tens of seconds (or more) using FEM [19] and accelerated BEM [11]. Moreover, using the fastest FEM/BEM code, the computational requirement of these direct approaches makes it almost impossible to solve for so many experimental simulation runs and find optimum coil placements across individuals.

Recently, there have been attempts to develop fast E-field solvers with the aim to achieve real-time (or near) solutions. A deep-learning-based E-field solver was proposed for real-time E-field analysis [20]. This approach is able to determine visually accurate E-fields in 0.024 seconds. However, the errors are not as low as standard E-field solvers limiting its usability for coil placement optimization. Another group [21] developed a highly optimized BEM implementation that can deliver real-time E-field distributions on a realistic head model. However, the CPU resources of their method scale with the number of tissue boundary triangles times the number of cortical samples times the number of coil model dipoles. Correspondingly, trade-offs between these three variables must be made to achieve realtime performance. Our method, in contrast, only scales with the number of E-field evaluation points enabling the use of high head model resolution when evaluating the E-fields. More recently, a magnetic profile approach was developed for real-time E-field evaluation for head models [22]. While this method seems promising, no results beyond 120,000 triangles were shown and typical FEM meshes have approximately 1 Million boundary triangles. Furthermore, their method required 1,500 – 3,000 simulations to generate the Huygens equivalent surface current that replicates the E-field to 5 – 15% relative residual error. In contrast, here we adopt a probabilistic matrix approach that enables us to achieve below 2% error with 220 simulations.

We recently introduced the ACA-ADM for E-field analysis and showed that it could determine the E-fields rapidly [23]. The method relied on using adaptive cross approximations (ACA) to compress a matrix containing the E-field distribution for a large number of scalp coil placements. Specifically, each column of the matrix defines the E-field throughout the whole brain for fixed coil placement. Since coils reside outside of the head and the brain inside it, the singular values of the matrix will decay exponentially [24, 25, 26]. Not unlike ACA-ADM we also approximate the mapping between coil placement and TMS-induced E-field into a matrix. However, we adopt a probabilistic matrix decomposition (PMD) to compress the matrix. The benefits of the probabilistic approach are twofold. First, it requires almost 2 times fewer simulations than ACA, and our numerical experiments indicate that it is near optimal. For example, for results in a spherical head model, the ACA and PMD required approximately 2.0 and 1.4 times the number of simulations, respectively, relative to the optimal compression using SVD. Second, unlike the ACA-ADM, we can infer the accuracy of the results from the PMD output, thereby being more robust.

We show simulation examples illustrating applications of our solver to find group-level optimum coil placements. First, we co-register the MRIs to an average subject template and use EEG 10/10 coordinates to specify cortical locations across individuals. Second, we use individualized EEG electrode locations as fiduciaries to generate common scalp coordinate systems across them. The mean and standard deviation of the E-field results across subjects are then shown in a human cortex template. This allows us to conduct ROI analysis across subjects and determine coil position that maximizes the E-field and minimizes its variance across subjects.

The rest of the paper is organized as follows. First, we describe the proposed approach of generating a standardized EEG-coordinate system across subjects. Then, we describe the low-rank matrix approximation technique for predicting the E-field for a large number coil placements over the scalp. This method is then bench-marked in terms of accuracy by comparing it with the FEM results. Furthermore, a detailed parametric study is conducted to determine the performance of the proposed method in terms of accuracy, speed, and chosen coil placement locations. All of these are done using a virtual population of five compartment head models of 200 subjects. Finally, our solver is used for population-level ROI analysis.

The software implementation of the proposed method can be found on GitHub (**Link to be available after publication**). The code can be used in the MATLAB environment. (Note: for the range of frequencies of TMS pulses (1 - 10 kHz), quasi-stationary assumptions are valid. As such, we assume that temporal and spatial variations are separable and the temporal variation is suppressed in what follows).

## 2. Methodology

Our virtual population-level E-field dosimetry estimation workflow is similar to the ones used by standard fMRI and TMS group-level analysis studies as illustrated in Fig. 1. Head models of each subject are generated from structural MRI images using SimNIBS’s ‘mri2mesh’ module [27]. A mapping between group-level scalp coordinates and individualized coordinates are defined using EEG 10/10 electrode positions as reference location (details in sections 2.5, 2.6 and 2.7). Group-level coil placements are defined on the scalp and mapped to each individual subject. The E-field is determined for each subject using our fast E-field solution method (details in section 2.1). The brain E-field results are co-registered for group-level analysis using Freesurfer’s ‘FsAverage’ human brain template and group-level E-field statistics are extracted [28] (details in section 3.6).

**Figure 1:**
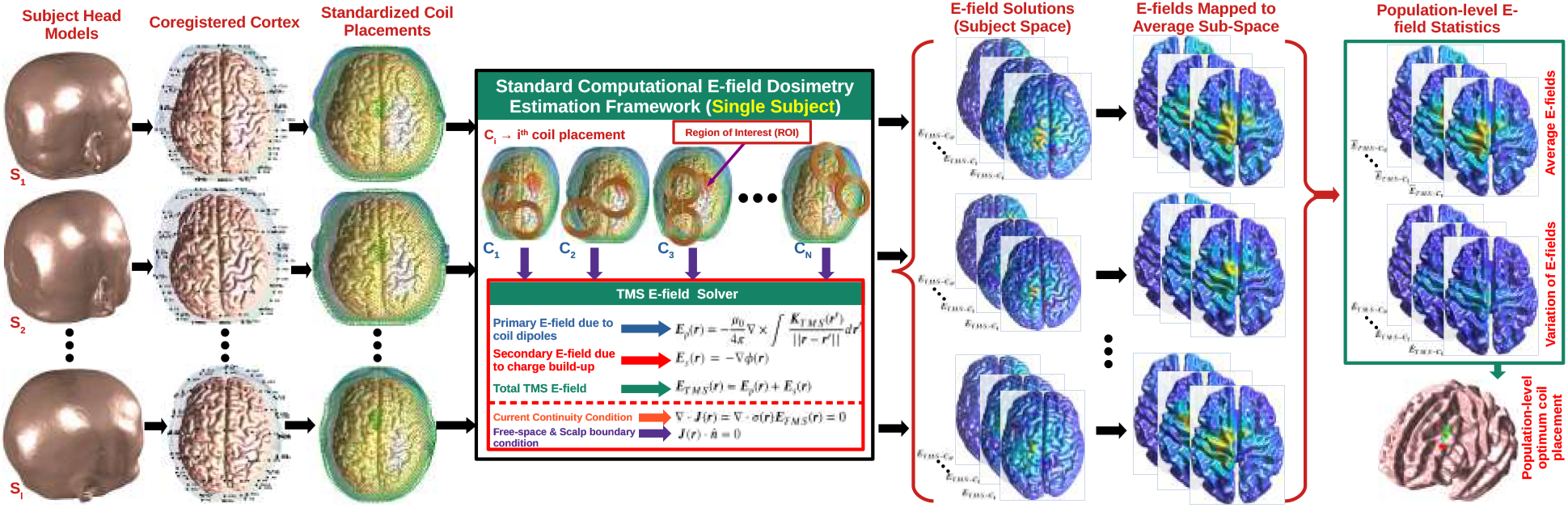
Population-level TMS E-field estimation and optimization workflow. The subjects’ head models are coregistered based on an EEG 10/10 electrode system and a set of group-level standardized coil placements are defined across subjects by considering the structural variation of the brains. The E-field is determined for all coil placement of every subject using a TMS E-field solver (FEM) and the fields are mapped to an average subspace (template). The statistical inferences (average and variation of E-fields) of these mapped E-fields are used to determine the optimum coil placement for an ROI across subjects.

The computation of the E-field for a large number of coil placements is a computationally intractable process as it requires a high number of simulations scaling as the product of coil placements and the number of subjects. The main contribution of this paper is to reduce this computational cost to the number of subjects times a fixed number of simulations, that is, virtually independent of the number of coil placements, and head mesh resolution via a probabilistic matrix approximation algorithm (details in section 2.2 and 2.3).

### 2.1. Computational E-field dosimetry

In this section, we describe the standard FEM and auxiliary dipole method (ADM) solvers used in this study. Computational E-field dosimetry requires a head model of the subject, a coil model, and relative placement of the coil over the subject’s scalp. The coil is modeled by a magnetic current and it generates a primary E-field that can be computed via Biot-Savart Law as

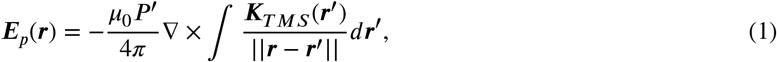

where ***μ***_0_ = 4***π*** × 10^−7^ *Am*^−2^ is the magnetic permeability of free-space, ***P’*** is the peak-time derivative of the current, ***K_TMS_***(***r’***) is the magnetic dipole moment at location ***r’***. (Note: here we have assumed that the E-fields exhibit a temporal dependence that is equal to the temporal derivative of the current pulse waveform, and can be rescaled to determine the temporal variation of the E-field.)

The primary E-field is distorted by a secondary E-field in the head that is generated by charge build-up at tissue interfaces. Furthermore, the secondary E-field is conservative (i.e. ***E**_s_*(***r***) = −∇*ϕ*(***r***), where *ϕ*(***r***) is the scalar potential) and the total E-field is ***E_TMS_***(***r***) = ***E**_p_*(***r***) + ***E**_s_*(***r***). The standard FEM solver uses eq. 1 to determine the primary E-field. From the primary E-field, the secondary E-field is then determined using current continuity. Current continuity dictates that the conduction currents in the head must be purely solenoidal (i.e. ∇ · ***J***(***r***) = ∇ · ***σ***(***r***)***E**_TMS_*(***r***) = 0, where ***σ***(***r***) is the conductivity of the head tissue). This results in the following Poisson equation for the scalar potential with a primary E-field-related forcing term

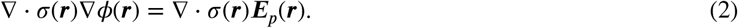

Additionally, free space is insulating at the frequencies of TMS, which results in 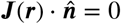 at the scalp surface. To solve eq. 2, the head is partitioned into a mesh consisting of tetrahedrons and the scalar potential is approximated via a 1st order nodal element expansion. Then, weak forms of the equations are derived using a Galerkin procedure. Finally, the resulting linear system of equations is solved for the coefficients of the expansion.

The standard FEM solver described above enables the computation of the E-field distribution in the head due to any fixed coil placement. In contrast, the auxiliary dipole method (ADM) enables the rapid computation of the E-field for a fixed location and direction of a large number of coil placements. The details of ADM can be found in [29] and next we provide a brief overview.

ADM computes the average E-field along a predetermined direction 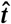 in an ROI. First, an auxiliary electric current source, flowing along the 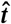 direction with constant intensity and having support on the ROI, is defined (i.e. 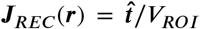, where *V_ROI_* is the volume of the ROI and ***r*** is in the ROI). The Electromagnetic Reciprocity principle is then used to relate the H-field generated by the auxiliary source (***H**_REC_*(***r***)) to the TMS-induced average E-field in the ROI

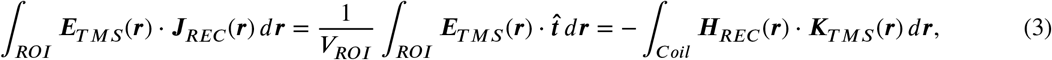

where ***Coil*** and ***ROI*** indicate the support of the coil magnetic dipole currents and ROI, respectively.

To compute ***H**_REC_*(***r***) outside the head, we first solve for ***E**_REC_*(***r***) by applying current continuity in the head. In other words, ∇ · *σ*(***r***)***E**_REC_*(***r***) = −∇ · *σ*(***r***)∇*ϕ_REC_*(***r***) = ∇ · ***J**_REC_*(***r***). This equation is solved using the standard FEM and the magnetic field outside the head is computed via Biot-Savart law as

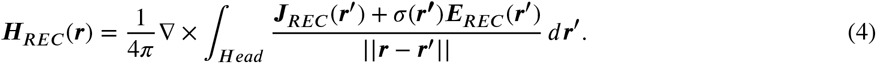

In this study, all E-field solutions are obtained using our 1st order FEM in-house solver [12] and ADM solver [29], respectively.

### 2.2. Matrix Approximation Formalism

During the population level analysis, we typically choose *N* distinct group-level coil positions. Each position is associated with 360 distinct coil placements by rotating the coil *1^°^* apart along its tangent plane on the scalp. Hence there are 360*N* distinct coil placements. For each coil placement we store the average E-field in each of the brain tetrahedrons. The results are compiled into a matrix ***A*** as

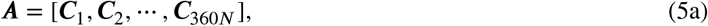

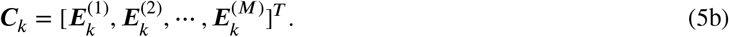

Here each column ***C**_k_* (*k* ∈ {1, 2,…, 360*N*}) contains E-field samples 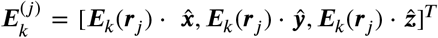, where ***r**_j_* is the center of one of the *M* brain tetrahedrons and *j* ∈ {1, 2,…, *M*}.

The matrix ***A*** can be computed column-wise using 360*N* standard FEM or row-wise using 3*M* ADM simulations. Assuming 2500 coil placements and 1.5 Million brain tetrahedrons, this would take approximately 5 years per subject using our in-house solver. To avoid this computational bottleneck, we previously used the adaptive cross-approximation method (ACA) to construct this matrix by interpolating results from judiciously chosen columns and rows [23]. However, the ACA-based compression still required 298 – 384 simulations (suggested 400) to compute the matrix to 2% accuracy and did not provide error estimates. Therefore, we have developed a more robust probabilistic matrix approximation approach described in the next section.

### 2.3. Low-Rank Matrix Approximation Algorithm

In our previous work, we showed that ***A*** is of low rank and compressible [23]. To approximate it, here we follow a probabilistic matrix decomposition approach designed to compress low-rank matrices [5]. The procedure to generate the probabilistic matrix decomposition involves left and right multiplication with the matrix ***A*** and is given in Algorithm 1. The method approximates the largest *r* singular values ***Ŝ***. Furthermore, it generates a basis for the largest *r* left and right singular vectors ***Q***, and ***B***, respectively.

#### Algorithm 1 Probabilitic Low-Rank Matrix Approximation

Given a matrix ***A***_3*M*×360*N*_ and an integer *r* (*r* = PMD approximation rank):

1. Construct a Gaussian random matrix **Ω**_360*N*×*r*_
2. Construct the matrix, ***Y***_3*M*×*r*_ = ***A*Ω**
3. Construct the matrix, ***Q***_3*M*×*r*_ from QR factorization of ***Y***, i.e. ***Y*** = ***QR***. The orthonormal columns of ***Q*** forms a basis set for the range of ***Y*** as well as for the range of ***A***
4. Construct a matrix ***B***_*r*×360*N*_ = ***Q**^T^**A***

In steps 1-2, an approximate basis for the space spanned by the first *r* left singular vectors of ***A*** is generated by multiplying a 360*N* × *r* matrix consisting of normally distributed entries (i.e. **Ω**) with it. In steps 3-4 the basis is converted to an orthonormal one (i.e. ***Q***) and the orthonormal basis is used to construct a basis for the first *r* right singular vectors by multiplying it on the right by ***A*** (i.e. ***Q^T^ A***). This process works because of two reasons. First, the columns of **Ω** have identical norms in expectation and are equally likely to point along any direction in the vector space. Second, the multiplication with the matrix acts to amplify vectors along the highest singular vector directions and suppress others, thereby, naturally generating a basis for the largest left and right singular vector spaces.

The standard FEM and ADM only enable the computation of columns and rows, respectively. As such, we do not have an efficient way to conduct the left and right multiplications with ***A***. To do the left multiplications efficiently, we linearly combine coil configurations and run a single FEM simulation as shown in the second row of Fig. 2. To do the right multiplications with ***A***, an ADM is run with a linear combination of auxiliary sources as shown in the fourth row of Fig. 2. The above procedures allow us to multiply with ***A*** on the right and left by a single standard FEM simulation and ADM simulation, respectively. This makes our approach require only a total of 2*r* simulation run-time plus a relatively small overhead introduced by the QR and SVD algorithm.

**Figure 2:**
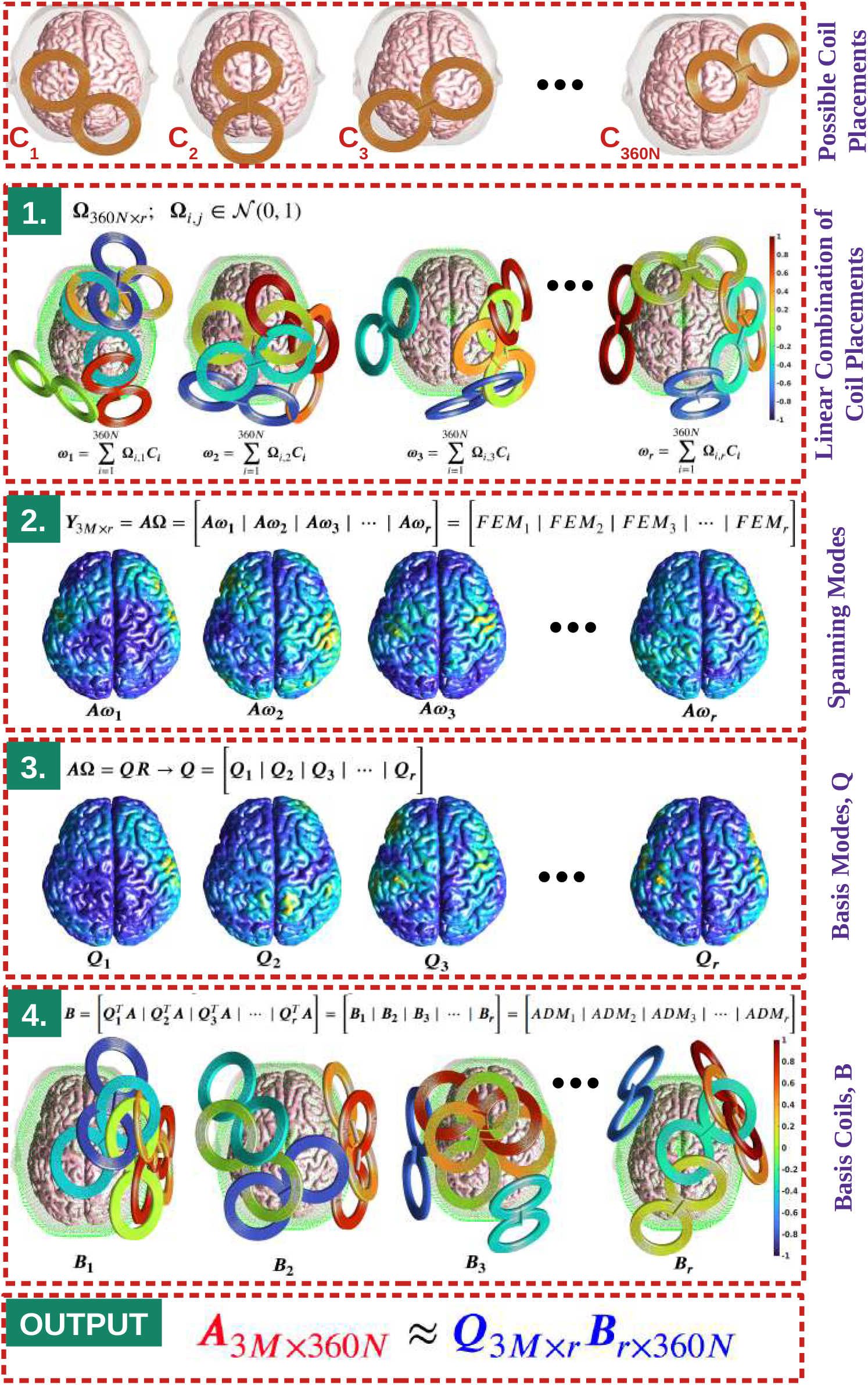
Visual Representation of the PMD algorithm. ‘Basis Modes’ and ‘Basis Coils’ represent the basis set for the column space and the row space of ***A*** matrix, respectively.

### 2.4. Spherical head validation study

For validation, we considered a spherical head scenario. The spherical head consists of one homogeneous compartment of radius *r_scalp_* = 8.5 cm and the coil consists of a single magnetic dipole oriented along the 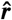 direction. We evaluate the E-fields in the head on a spherical surface of radius *r_obs_* = 7.0 cm [Fig. 5A-B]. Observation points are chosen as the nodes of a five times refined icosphere as shown in Fig. 5A-B. Additionally, coil placements are chosen on a spherical cap of radius *r_coil_* = 9.0 cm and cap height of 0.5 cm as shown in Fig. 5A-B. Analytical formulas are used to determine the E-field for each coil placement and magnetic dipole placement [30]. The resulting matrix is of size 30726 × 2553. To validate our method, we constructed this full matrix and compressed it using singular value decomposition, PMD, and ACA.

**Figure 3:**
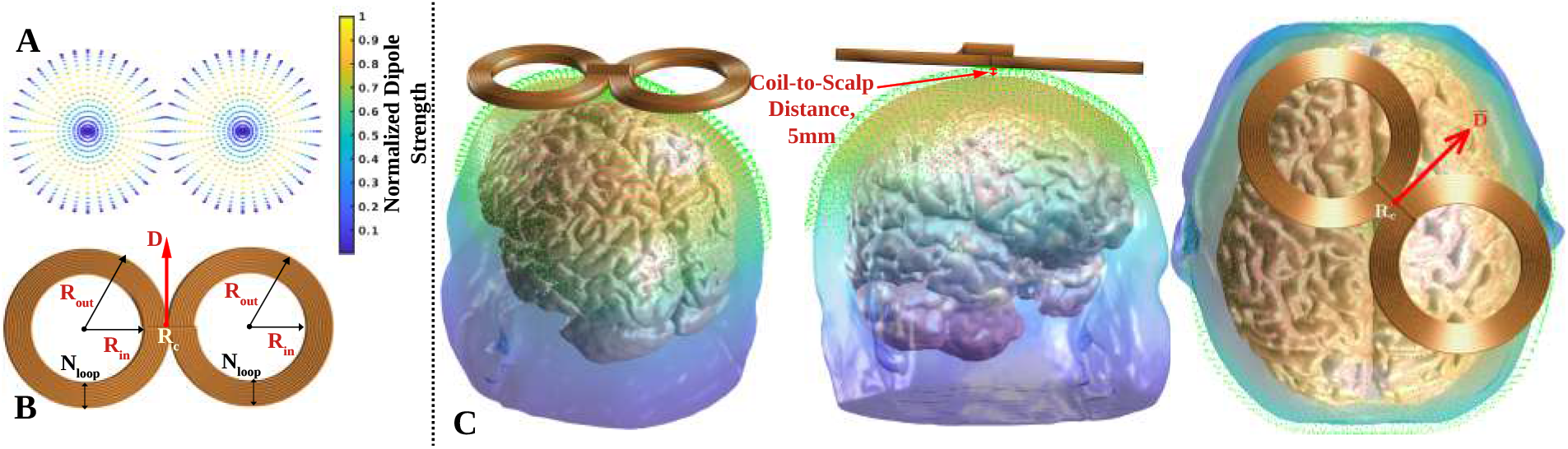
TMS figure-8 coil and placement. (A) Dipole model of the TMS coil. (B) The center of the coil (***R**_c_*) denotes the coil position and the red arrowhead denotes the coil direction (or, orientation), 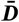. (C) Placement of TMS coil over the scalp, tangential to the local scalp region.

**Figure 4:**
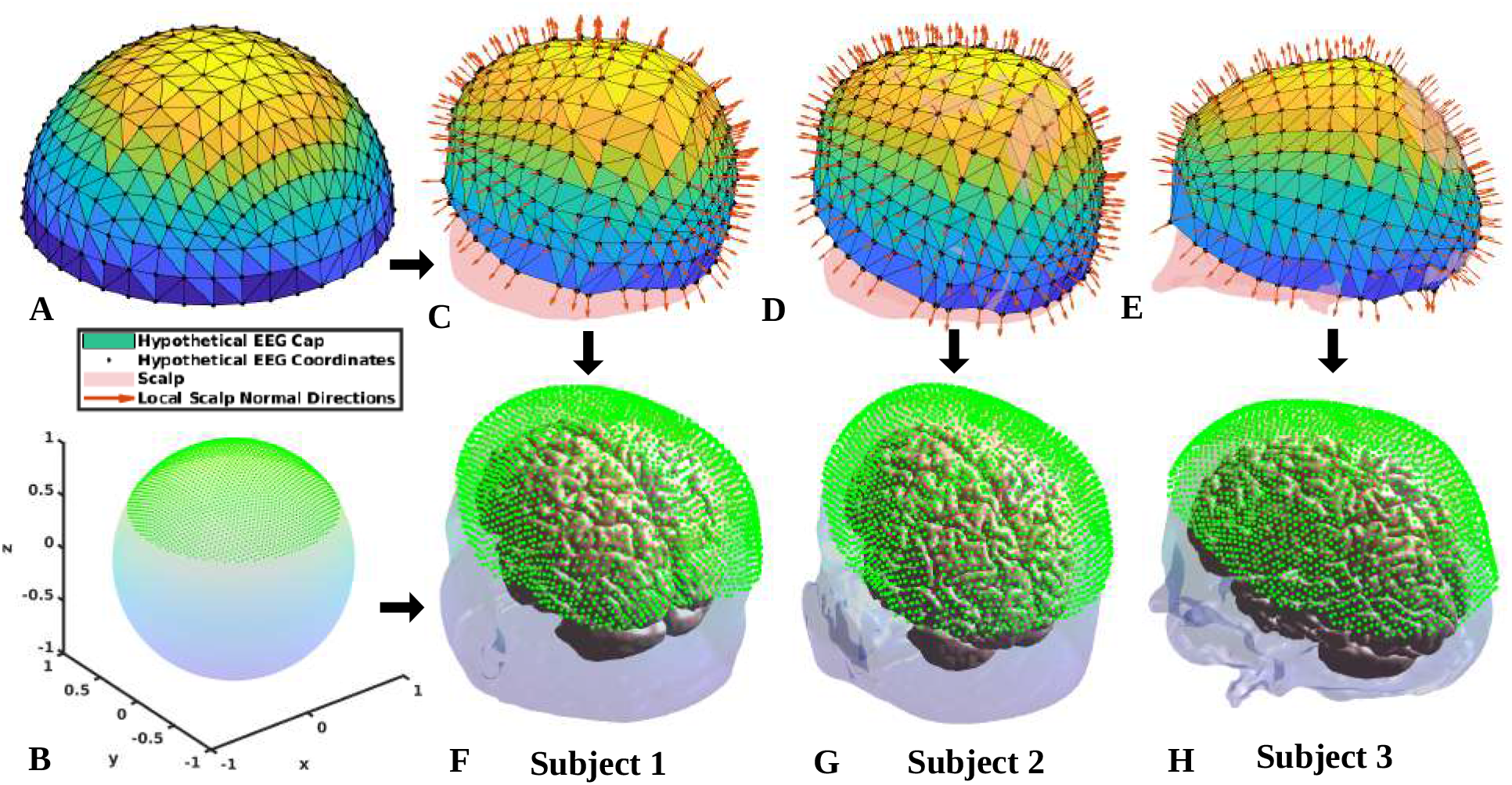
Construction of standard EEG coordinate system across multiple subjects. A standard EEG system is formed over a unit sphere (A). Next, uniformly distributed points are sampled from a unit icosphere model with an angular polar patch size of 60° (B). With the angular locations of the standard EEG system, an individualized EEG coordinate is constructed for each subject (C-E). Next, the sampled points from a unit sphere in (B) are projected onto the standardized EEG plane of each subject (F-H).

**Figure 5:**
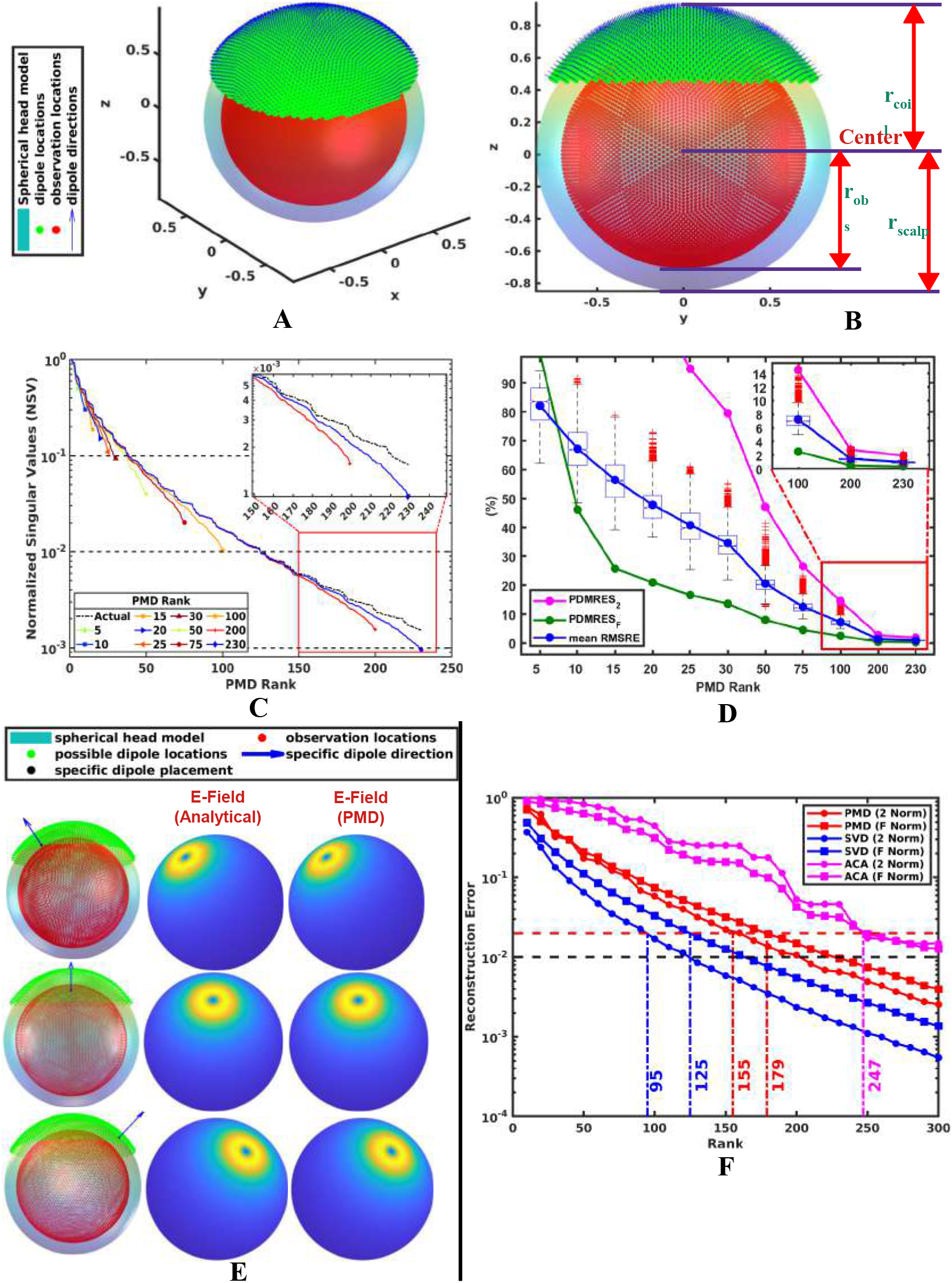
Compression of the coil position to E-field in a spherical head matrix using SVD, PMD, and ACA. (A-B) The spherical head model along with magnetic dipole placement centers and E-field sample points. (C) Comparison of actual vs PMD-approximated singular values for various PMD ranks. (D) ***RMSRE*** as a function of rank. (The distribution of ***RMSRE*** is calculated from 1000 random magnetic dipole placements.) The maximum ***RMSRE*** is below 2% at rank, *r* = 230. ***PMDRES***_2_ and ***PMDRES**_F_* act as the upper bound and the lower bound for the ***RMSRE***. (E) Comparison between the E-field solutions from the analytical model and PMD algorithm for random dipole placements. (F) Minimum ranks required by the SVD, PMD, and ACA algorithms to achieve a 2% Frobenius norm and 2-matrix norm after compression.

**Figure 6:**
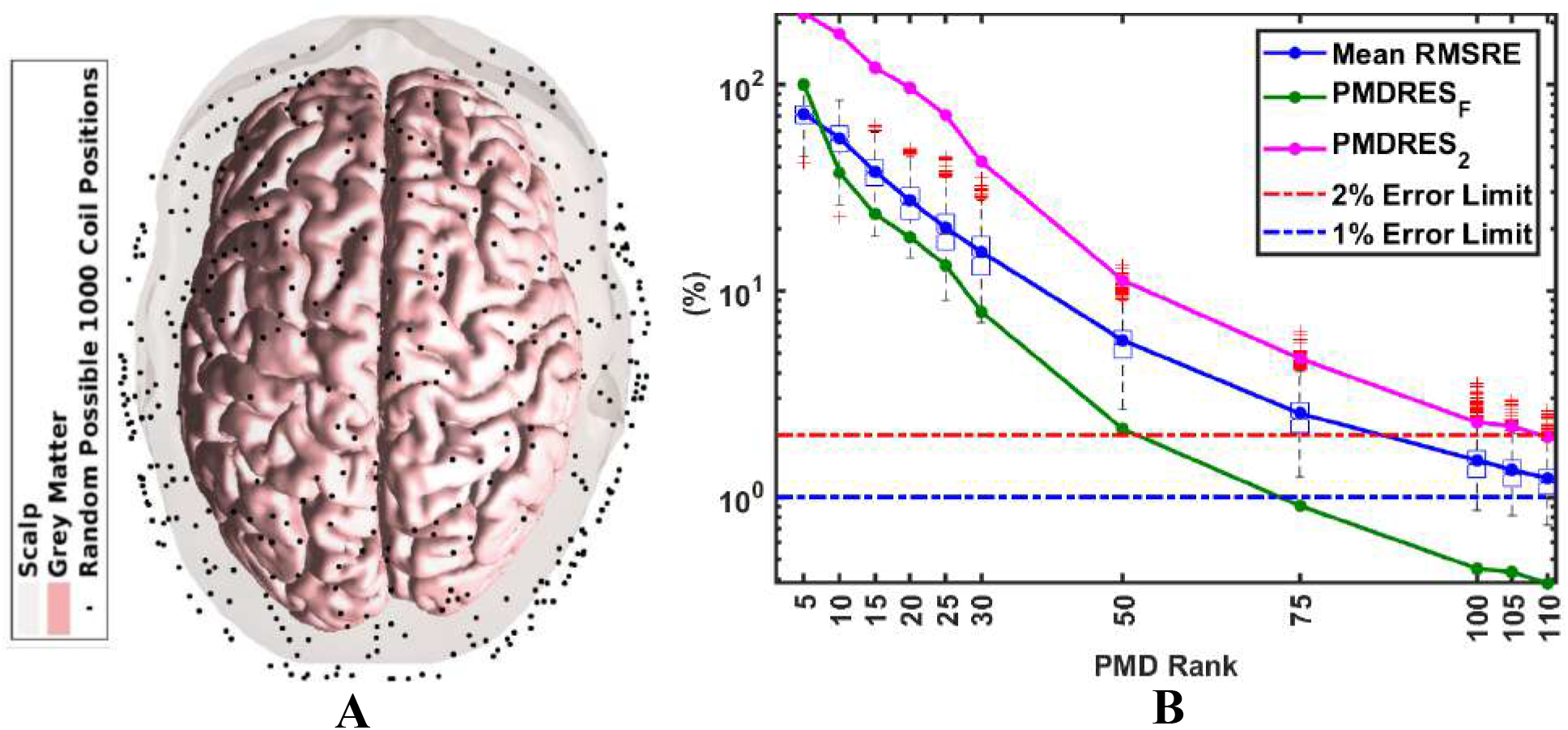
(A) The 1000 Randomly chosen coil positions over the scalp for a single subject. (B) ***RMSRE*** as a function of PMD rank. The box and whisker plots show the distribution of ***RMSRE*** of E-field over 1000 coil positions and the solid blue line represents the mean ***RMSRE*** at any rank. For the rank of 110, the mean value of ***RMSRE*** is approximately 1%, whereas the maximum ***RMSRE*** is below 2.5%. ***PMDRES***_2_ metric acts as nearly an upper bound for the ***RMSRE*** and ***PMDRES**_F_* is a lower bound for the error.

**Figure 7:**
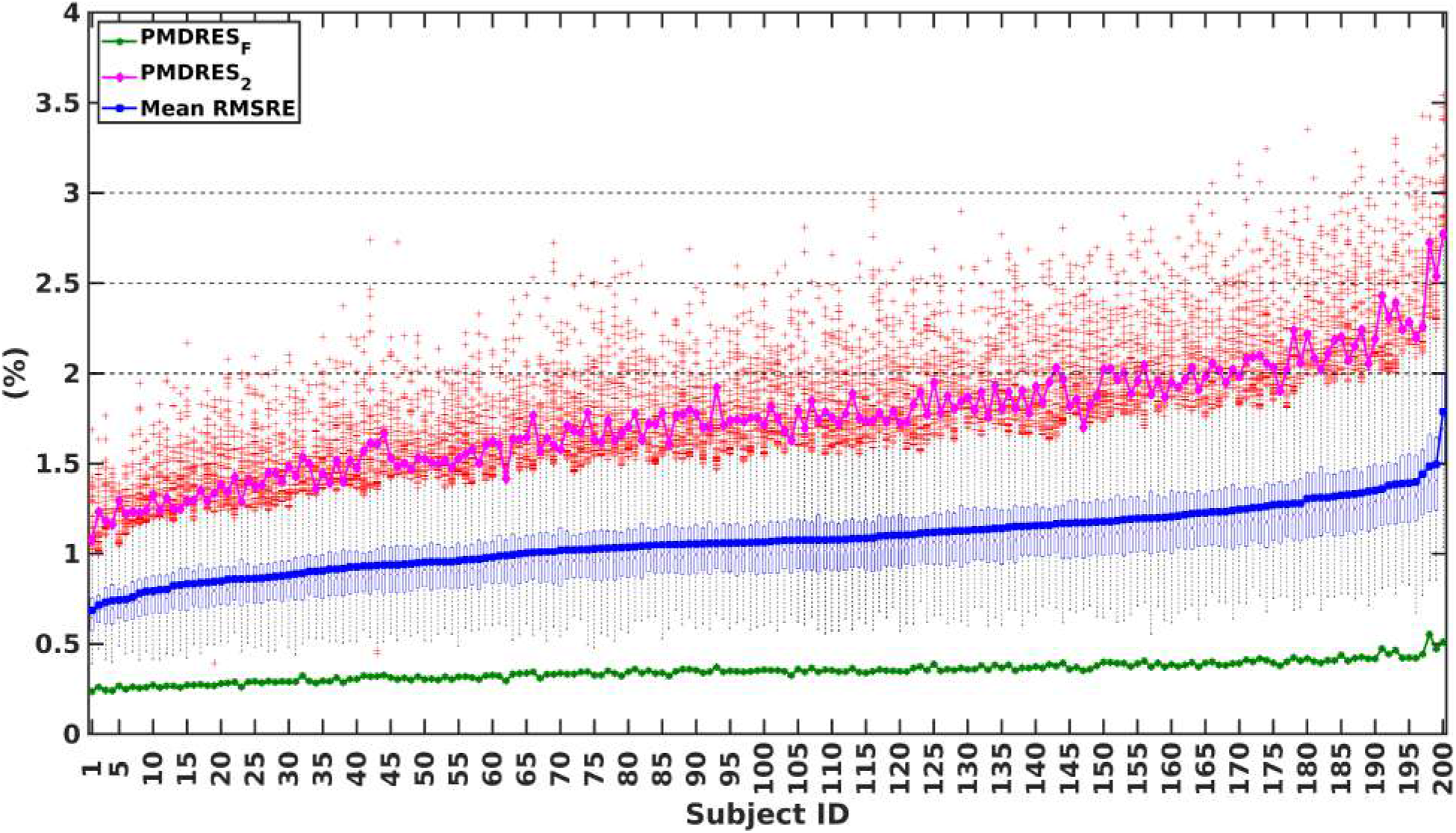
***RMSRE*** and PMD error metrics across 200 subjects when the PMD rank (*r*) is fixed at 110. The mean value of ***RMSRE*** is around 1% across all subjects. For all subjects, the maximum ***RMSRE*** is within 3.55%. (For every subject the distribution of ***RMSRE*** is calculated across 1,000 random simulations.)

**Figure 8:**
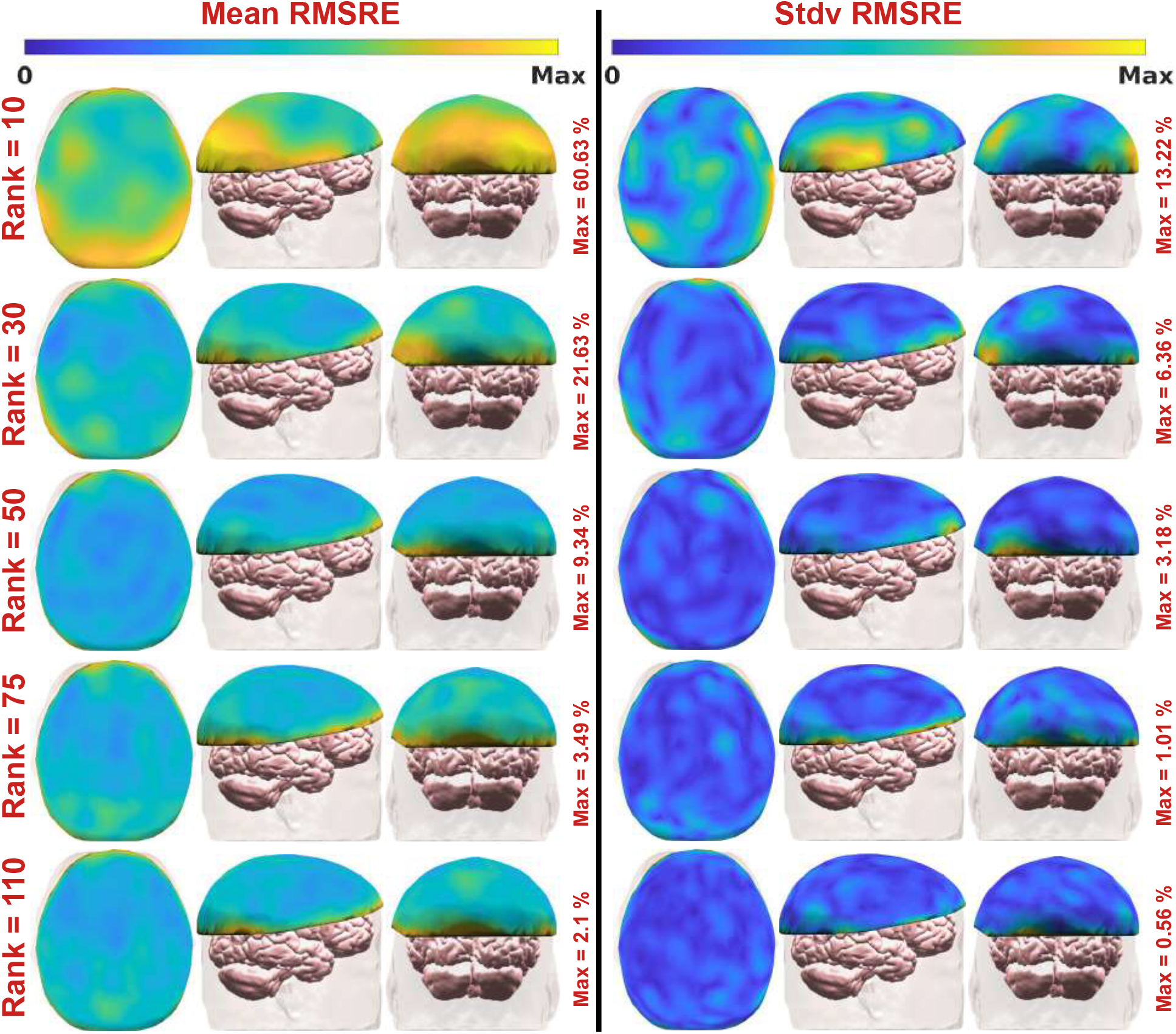
Illustration of ***RMSRE*** as a function of coil position. The respective mean and standard deviation (Stdv) of the ***RMSRE*** are shown for increasing PMD rank top to bottom. The first three and last three columns are the axial, sagittal, and coronal views of the mean and Stdv, respectively. The higher ***RMSRE***_s_ are concentrated near the periphery of the standardized EEG cap due to the abrupt truncation of the standardized EEG mesh. In each case the Stdv as a function of angle is small indicating low dependence of RMSRE to coil orientation.

### 2.5. Virtual Head Model Generation and co-registration

To test the proposed PMD algorithm and do population-level analysis, we developed a virtual head model (VHM) database from Magnetic Resonance Images (MRI) of 200 subjects. The demographic information of the database is listed in Table 1. The MRI scans are collected from three distinct studies [31, 32, 33]. For each subject, the T1 and T2 weighted sequences are used to generate a VHM. The VHM is generated by segmenting and generating a 5-compartment tetrahedral mesh using the ‘mri2mesh’ tool of SimNIBS [19]. The ‘mri2mesh’ uses FSL’s [34] brain extraction tool to extract intracranial compartments and generate a CSF boundary mesh. This mesh is then extruded outward to generate the skull. The gray matter (GM) and white matter (WM) surfaces are obtained using Freesurfer [28]. All surface meshes are then merged and a volume mesh is generated using GMSH [35]. The output meshes of each subject have approximately 4.5 Million tetrahedra and consist of five homogeneous compartments GM, WM, cerebrospinal fluid (CSF), skull, and scalp. Each virtual head is converted to a volume conductor model (VCM) by assigning each region its respective conductivity provided in Table 2 and are chosen from values given in [36]. The average time needed for the generation of each VCM is approximately 25 hours. For our analysis, we only consider the E-fields induced in the GM and WM regions (i.e. the brain) and the corresponding total number of brain tetrahedrons (*M*) is approximately 1.5 Million. To conduct our group-level analysis, we use Freesurfer’s ‘FsAverage’ template to map the E-field solutions from every subject.

**Table 1.** Database Demographic Information

| Summary | Count |
| --- | --- |
| Number of Subjects | 200 |
| Virtual Head Models (VCMs) | 200 |
| Number of Male Subjects | 102 |
| Number of Female Subjects | 98 |
| Healthy VCMs | 200 |
| Number of MRI Sessions | 200 |
| Subject Age Range | 18-95 Years |

**Table 2.** Standard Tissue Conductivities used for VCMs

| Tissue Type | Conductivity Value (S/m) |
| --- | --- |
| White Matter | 0.126 |
| Gray matter | 0.275 |
| Cerebrospinal Fluid | 1.654 |
| Skull | 0.01 |
| Scalp | 0.465 |

### 2.6. TMS Coil Model

Unless stated otherwise in all simulation studies, the coil is chosen to be a 70mm Figure-8 coil (#31 in [37]). The inner and outer radii (***R**_in_*, ***R**_out_*) of the coil loops are 28*mm* and 43.5*mm*, respectively [Fig. 3B] and it has 2 layers and 9 loops (*N_loop_*) [Fig. 3A]. The model of the coil consists of 2880 magnetic dipoles as depicted in Fig. 3A. This coil model has been used in our previous studies and shown to generate primary E-fields with errors ≤ 1% when compared with higher-resolution dipole models [12]. The coil is always assumed to have a *d**I**/di* = 6.7 × 10^7^A/s.

In this study the coil is placed tangent to the scalp directly below its center. The coil positioning relative to the head is identified by the location of its bottom and center, and an orientation angle ***ϕ’***. Specifically, we always assume that the coil position is always 5*mm* directly above the scalp and an orientation angle ***ϕ’*** defines a rotation relative to the anterior to the posterior axis. The 5 mm displacement between the coil wire and scalp accounts for coil casing and hair thickness. As in Fig. 3B, the coil is placed at a position (***r’***) over the scalp in such a way that the coil center (***R**_c_*) is overlapping with (***r’***) and the coil direction 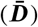 is overlapped with the orientation (***ϕ’***). Furthermore, we consider integer angle orientations 1*^°^* – 360*°* (1*°* apart) for every coil position.

We apply minor Gaussian smoothing to the scalp mesh before computing normals to avoid stair casing artifacts from the voxelized MRI image. To define normals anywhere on the scalp, we first find the scalp node normals by averaging the normals of the surrounding triangular facets [38]. Then, we interpolate the normals at each node to any scalp position by using standard 1st order nodal elements. Note: even though coil placement is assumed tangential to the scalp, the proposed approach for determining the E-field using the probabilistic matrix decomposition technique is equally effective for arbitrary coil placement (i.e. including a coil placement with feasible center coordinate location, yaw, pitch, and roll directions).

### 2.7. Standardized EEG Coordinate System

Like in clinical applications of TMS to select consistent coil placement across subjects, we define coil positions in terms of standard EEG electrode placements. First, we generate a Delaunay triangle mesh from nodes on the hemispherical coordinate angles of the EEG 10/10 electrodes. The resulting mesh is barycentrically refined twice that results in the mesh shown in [Fig. 4A]. Second, we use fiducials at the Nasion (Nz), Left Preauricular point (LPA), Right Preauricular point (RPA), and Inion (Iz) to find individualized coordinate placements for each node angle [Fig. 4C-E]. Individualized meshes shown in Fig. 4C-E all share the same connectivity as the one in Fig. 4A. Third, dense samples on the spherical cap are chosen on the region with the polar angle between 0° and 60° as shown in Fig. 4B and individualized using meshes to generate coil placements shown in Fig. 4F-H.

For each coil position we consider 360 coil orientations (1° apart from each other). Unless stated otherwise, we choose *N* = 2655 coil positions over the scalp. This will result in 360*N* = 955, 800(≈ 1 *Million*) coil placements. This large number of placements is chosen to ensure that we can determine the coil position that maximizes the E-field induced at the targeted brain site and minimizes variability across subjects accurately.

### 2.8. Error Metrics

Theoretically expected error bounds for the *r* = *k* + *p* order PMD approximation of any matrix are given in [5] as

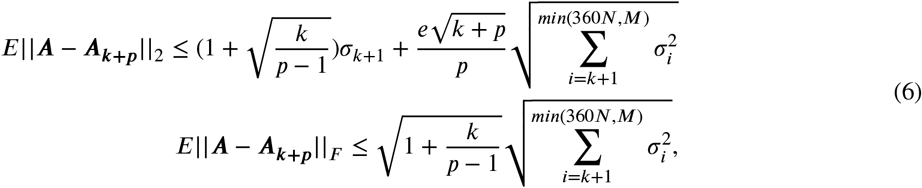

where *E*∥ · ∥_2_ and *E* ∥ · ∥_*F*_ are the 2-norm and Frobenius matrix norms, *e* is the exponential constant. Here *k* is the assumed rank of the matrix and *p* is an oversampling parameter. The actual rank used when running PMD is *r* = *k* + *p*. We use these error bounds to create error indicators for the algorithm. The singular values of the matrix ***A*** are known to decay exponentially from the theory of Laplace operators [26], and coil design literature [12]. As such, to create our error estimates, we assume singular values *σ*_*k*+2_ ≈ *σ*_*k*+3_ ≈ …*σ*_360*N*_ ≈ 0. This approximation results in the replacement of the sums in Eq. 6 with *σ*_*k*+1_. Furthermore, in Eq. 6, we replace the singular values with those approximated using PMD. Finally, we normalize the error bound with its respective matrix norm to attain the following probabilistic matrix decomposition relative error estimates (PMDRESs) in Eq. 7

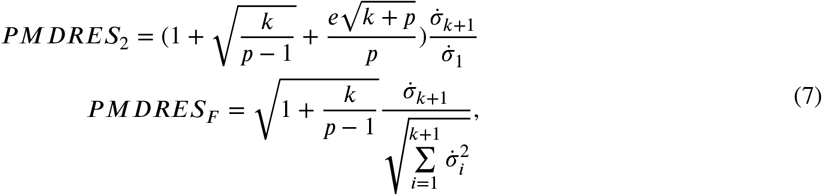

where 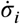 are PMD approximate singular values. Our results show that choosing *p* = 5 (i.e. neglecting the last 5 singular values) results in ***PMDRES**_F_* and ***PMDRES***_2_ values that provide a good estimate for the actual error.

The root mean squared relative error (***RMSRE***) between the FEM simulated E-field and the approximated E-field from the proposed PMD algorithm is calculated using equation 8 as

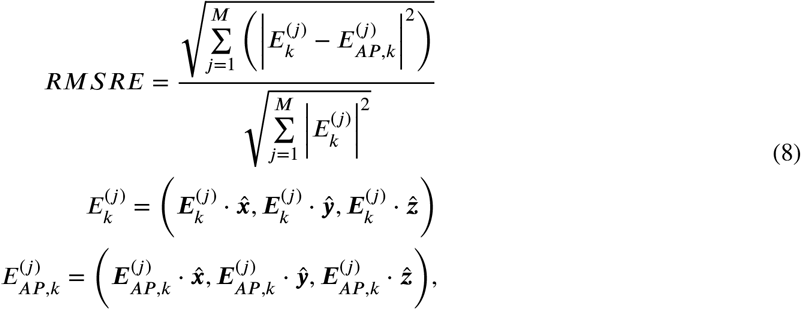

where, 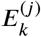 and 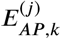 are the FEM generated E-field and the PMD approximated E-field, respectively, in the *j^th^* tetrahedron due to *k^th^* coil placement over the scalp. We also bench-marked our algorithm in terms of accuracy by using different similarity metrics such as Pearson Correlation Coefficient (PCC) [39], Cosine Similarity(CS) [39], Triangle Area Similarity(TS) [40], and Sector Area Similarity(SS) [40] which have been previously used to benchmark various brain stimulation E-field dosimetry tools [20, 21, 22, 7, 10, 41]. Details can be found in section S1 and Fig. S1. The errors for these metrics were all orders of magnitude smaller than 1%, as such, are only included in the supplementary material for reference only.

## 3. Results

### 3.1. Analytical Solution vs PMD over a Spherical Head Model

First we consider the compression of a coil placement to the E-field matrix for a sphere head model. The singular values of the ground-truth matrix along with the PMD compressed one are shown in Fig. 5C. The singular values decay exponentially [Fig. 5C]. The PMD algorithm produces good singular value estimates. As expected, the last few singular values are systematically slightly underestimated. Since the trends in singular values are maintained, the PMD provides adequate estimates for singular values that can be used in error tolerance calculations.

Fig. 5D shows the actual ***RMSRE***, ***PMDRES***_2_ and ***PMDRES**_F_* as a function of PMD rank. Overall, the ***RMSRE*** decreases with increasing PMD rank. In other words, PMD converges monotonically. The ***PMDRES***_2_ is higher than the actual expected value of the ***RMSRE***. The ***PMDRES**_F_* is lower than the actual expected value of the ***RMSRE***. Lowering the ***PMDRES***_2_ to 2.5% resulted in a maximum ***RMSRE*** of 2% for all coil placements. Fig. 5E shows some representative sphere results and they look visually identical. All of these results indicate that the PMD method can produce an accurate compression of the coil position to the E-field matrix.

Fig. 5F shows the 2-matrix norm and Frobenius matrix norm error achieved after matrix compression with SVD (i.e. optimal compression), PMD, and ACA. The PMD reconstruction error is much closer to the SVD error than the ACA error. For example, to achieve a 2% Frobenius error, the SVD, PMD, and ACA require ranks of 125, 179, and 247, respectively. In other words, the PMD achieved 1.38 times optimal compression, whereas the ACA was 1.98 times optimal compression. The PMD achieves near-optimal compression of the matrix while only requiring left and right multiplication with it.

### 3.2. Accuracy convergence of PMD as a function of rank

Here, we consider the suitability of the ***PMDRES*** metrics for predicting the actual accuracy of the matrix compression in an MRI-derived head model. Fig. 6 shows comparison results between FEM and PMD compressed results for 1000 randomly chosen coil placements [6A] on a single subject. The distribution of the ***RMSRE*** as a function of rank is shown in Fig. 6B along with the PMD metrics. The PMD metrics and ***RMSRE*** decrease with increasing rank. Furthermore, the ***PMDRES***_2_ nearly forms an upper bound for the actual error. For example, for a PMD rank of *r* = 110, the ***PMDRES***_2_ = 2% and the actual ***RMSRE*** for 1000 random simulations are below 2.5%. Furthermore, the mean of the ***RMSRE*** is near 1%.

Fig. 7 shows the ***RMSRE*** distribution for 200 subjects for a PMD rank of 110. Each distribution is calculated from 1000 coil position samples. We observe that for *r* = 110, all of the results had RMSRE errors below 3.54%, 99% of them are below 3% and 98% of them are within 2.5%. Furthermore, the mean ***RMSRE*** is near 1%. These results indicate that ***r*** = 110 is sufficient to ensure that the PMD compression does not result in significant additional numerical error than FEM. Furthermore, the PMDRES metric correlates well with the mean error indicating that it provides a reliable measure of the actual error. Several outliers are present in the simulation and they have errors about twice the mean ***RMSRE***. The coil positions that have the highest errors are the subject of the next subsection.

The results shown here involve a matrix that is 3***M*** × 360*N*, where ***M*** ≈ 1.5 × 10^6^ and 360*N* ≈ 10^6^. Furthermore, the matrix can be stored to 2% accuracy by storing only 110 column basis vectors and 110 row basis vectors, which is four orders of magnitude smaller than either matrix dimension and, correspondingly, we only need to store one < 0.01% of the matrix to generate the whole matrix. As a result, the compression in required storage is over 10^5^ times.

### 3.3. Error distribution as a function of coil placement and cortical location

Fig. 8 shows the ***RMSRE*** error as a function of coil position with coil orientation fixed. For each coil position, the mean and variation of the ***RMSRE*** are calculated for uniformly sampled coil orientation at 18 different angles from 1*°* to 180*°* (10*°* apart from each other). Nearly double ***RMSRE*** is observed on the boundary of the coil positioning mesh. As such, the increased error (i.e. outlier error cases in the previous section, in Fig. 7) is likely due to truncation effects. To increase the accuracy, the coil positioning grid can be padded beyond admissible coil positions. The above would enable the use of a lower rank in PMD compression. However, this was not explored further here, as the least accurate result already matches the FEM beyond its numerical accuracy [12].

Fig. 9 shows PMD and FEM predicted E-field distributions for exemplary simulation results. Visually both the FEM and PMD results appear identical. The ***RMSRE*** for each of the scenarios is 1.12%, 1.55%, 1.29% is 0.89%, 1.18%, and 1.23%, respectively. Furthermore, a maximum local error in the E-field prediction of 5% was observed across all results shown here. The results indicate that the PMD-predicted E-field distributions are accurate.

**Figure 9:**
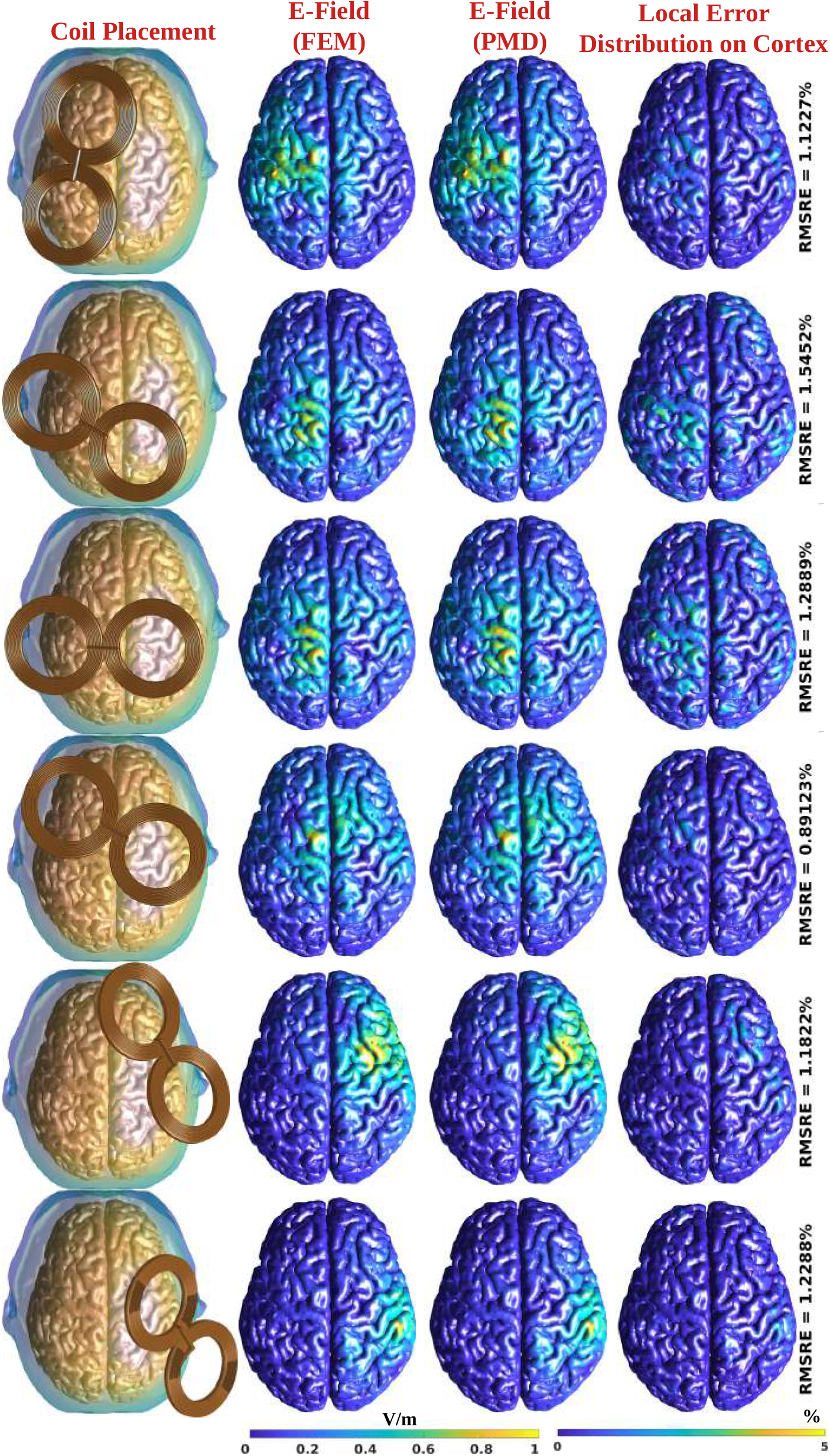
Illustration of the FEM-induced E-field (2^*nd*^ column) and PMD approximated E-field (3*^rd^* column) for randomly chosen coil placements (1^*st*^ column) over the cortical surface of a subject. The last column shows the local error distribution over the cortical surface along with the corresponding ***RMSRE*** value. The E-fields are represented in a normalized scale.

### 3.4. Coil position density and VCM resolution

We test the robustness of PMD with an increasing number of coil placements. For each test case, the locations of the coils are distributed uniformly on the same standardized EEG surface and their density is increased. Fig. 10 shows the PMD metrics and ***RMSRE*** as a function of rank and number of coil placement locations. ***PMDRES**_F_* and ***PMDRES***_2_ as a function of rank show negligible variation with an increasing number of coil positions [Fig. 10B]. Fig. 10A shows the ***RMSRE***, ***PMDRES**_F_*, and ***PMDRES***_2_ metrics for all test cases. Again, the results remain nearly identical as coil placement density is increased. The rank appears to be independent of coil grid placement density.

**Figure 10:**
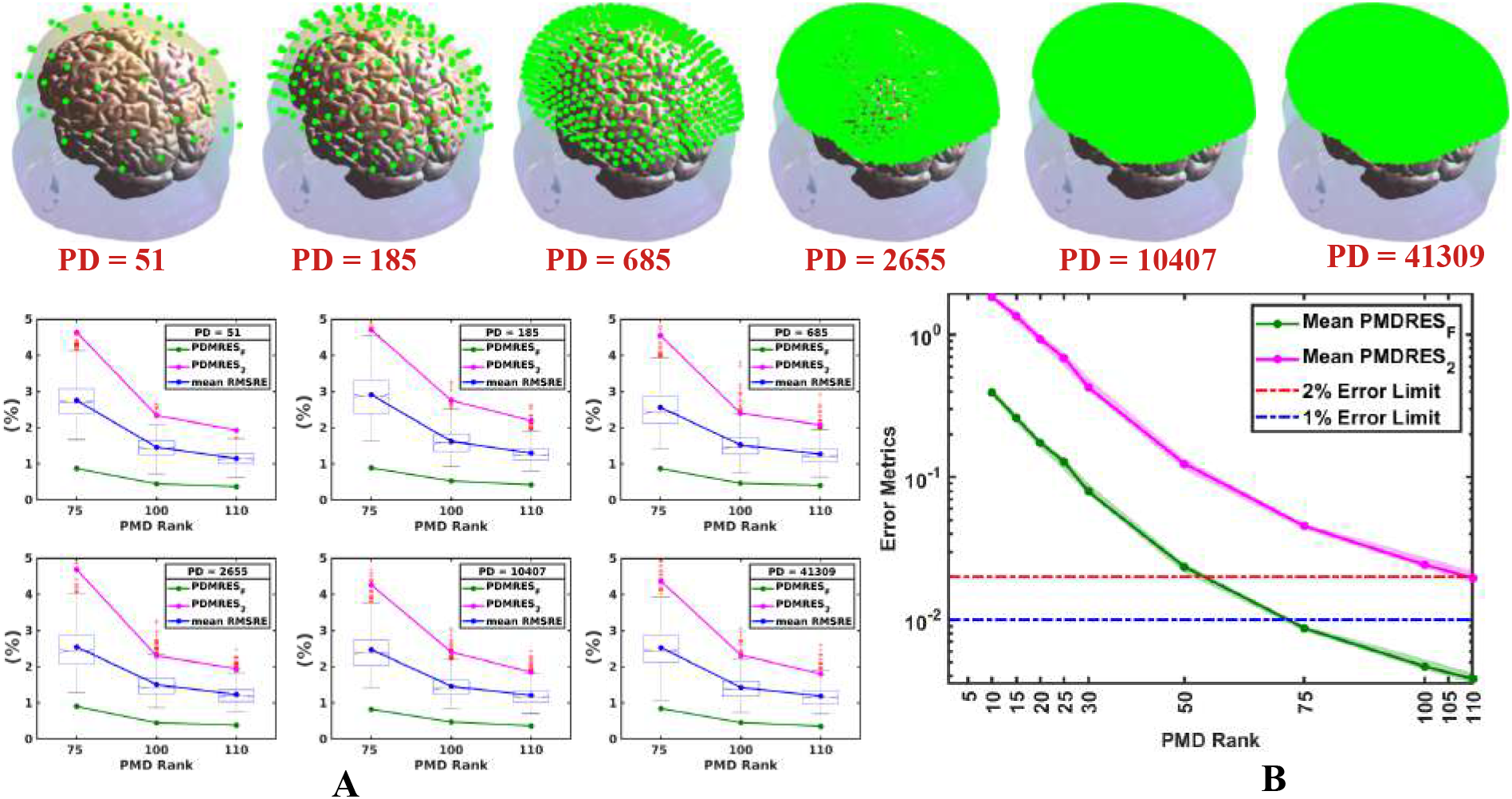
Variation of ***RMSRE***, ***PMDRES**_F_*, and ***PMDRES***_2_ with respect to rank as a function of point density on the standardized EEG coordinate system. The top row shows the standardized EEG points with increasing point density (from left to right). (A) The ***RMSRE*** and PMD error metrics for higher ranks (*r* ∈ {75,100,110}). For each scenario, the distribution of ***RMSRE*** is calculated across 1,000 random simulations. (B) The relative variation of ***PMDRES**_F_* and ***PMDRES***_2_ with respect to rank for different point density. Both of ***PMDRES**_F_* and ***PMDRES***_2_ are almost absolutely constant as a function of point density.

We consider the convergence of the PMD error metrics and ***RMSRE*** as a function of rank and head mesh density. A subject mesh consisting of 151,160 nodes and 877,106 tetrahedra is baricentrically refined twice to generate meshes with increasing resolution. The two additional meshes consist of 1, 184, 421 & 9, 405, 673 nodes and, 7, 016, 848 & 56,134,784 tetrahedra, respectively. Fig. 11 shows ***PMDRES**_F_*, ***PMDRES***_2_, and ***RMSRE*** as a function of rank for each of the meshes. The PMD metrics and ***RMSRE*** marginally decrease for a fixed rank as the resolution of the mesh is increased. The above results indicate that the rank necessary for accurate approximation of the FEM results is nearly independent of mesh density and is slightly smaller for higher resolution meshes.

**Figure 11:**
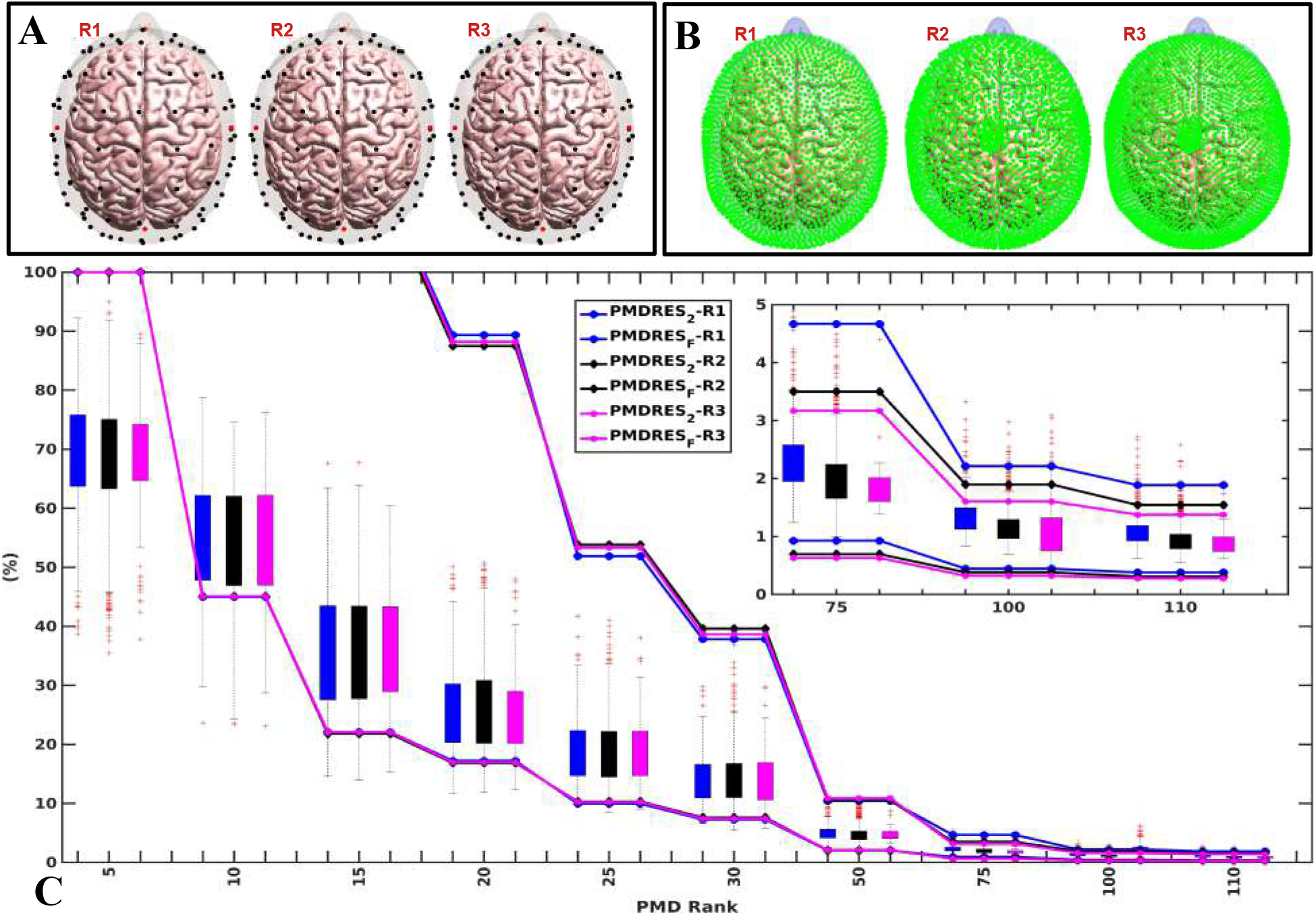
(A) Three different VCMs (***R***1, ***R***2, ***R***3) of the same subject with different resolutions with ***R***3 > ***R***2 > ***R***1. The head models are co-registered using an EEG 10/10 coordinate system. The red dots denote the four fiducial points (Nz, Iz, LPA, RPA). (B) The corresponding standardized EEG coordinates over the scalp. (C) ***RMSRE*** and PMD metrics (***PMDRES***_2_ and ***PMDRES**_F_*) as a function of PMD rank (*r*) for three different head resolutions. For PMD ranks, *r* ≥ 75, the ***RMSRE*** gets lower for increasing resolution. The PMD metrics act as error bounds.

### 3.5. Computational Run-time and Memory Requirements

Fig. 12A shows the average and 100^*th*^ percentile confidence intervals of the prepossessing time (set-up time) as a function of rank across 200 subjects. The set-up time for a single subject (assuming PMD rank, *r* = 110) ranges between 8.5 and 10.5 hours. This time is based on the in-house built FEM and ADM solver. Both of them require ≈ 170 seconds to solve for the E-field over the whole brain for single coil placement on a low-end cluster CPU (AMD Rome CPU, 2.0GHz). Furthermore, the time increases almost linearly with increasing rank. The variance in time required between subjects is due to slight differences in the FEM run times between subjects. Note that this time is significantly lower than the direct approach of calculating every column or every row of **A**. For example, with the same in-house developed code, the direct approach would have taken over 5 years for just a single subject.

**Figure 12:**
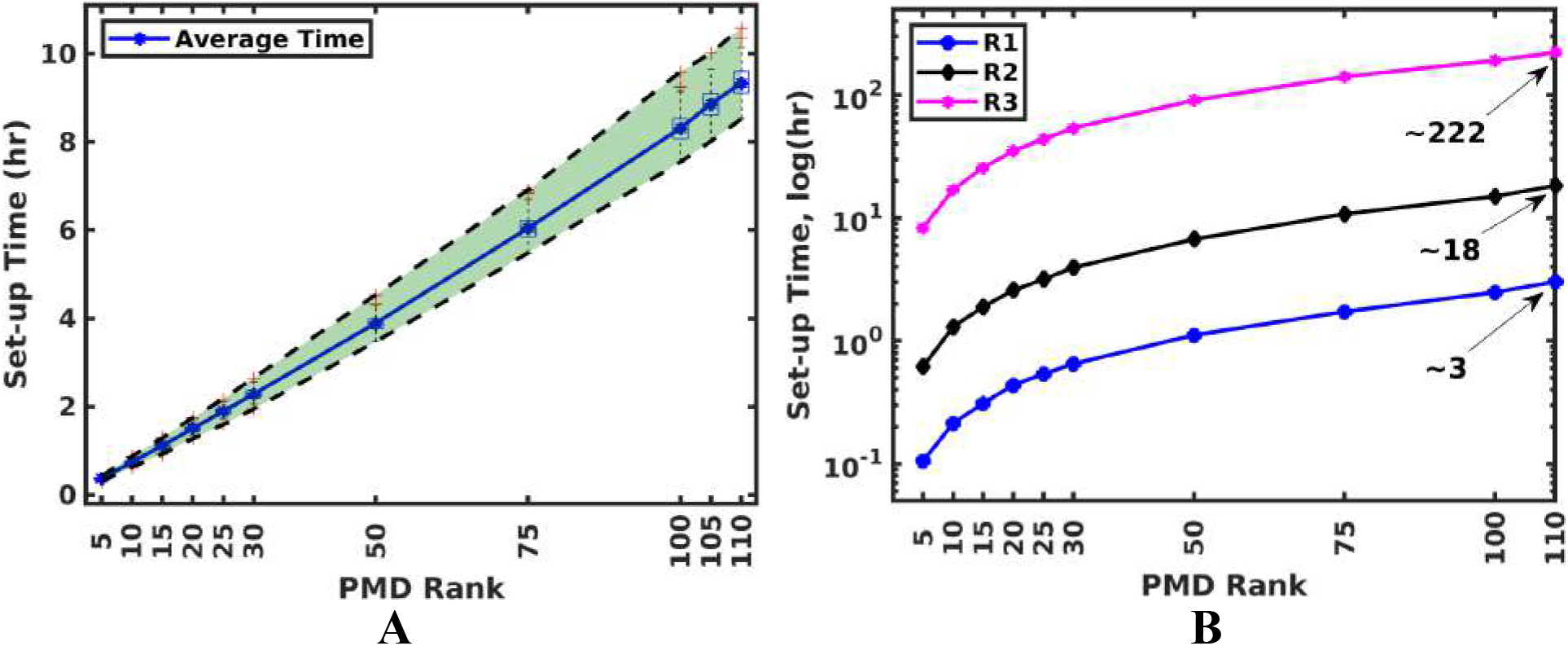
(A) Set-up time for a standard-resolution head model (≈ 1.5 Million tetrahedra). The 100% confidence interval is shown across 200 subjects. The set-up time increases almost linearly with respect to rank. The time is calculated on a low-end cluster CPU (AMD Rome CPU, 2.0GHz). At rank, *r* = 110, the set-up time is ≈ (9.56 ± 1.03) hours. (B) Set-up time for different resolutions of the tetrahedral mesh (***R***1 ≪ ***R***2 ≪ ***R***3). For every higher order of resolution in the tetrahedral mesh, the simulation time also increases by one order (3 hr to 18 hr and 18 hr to 222 hr).

Fig. 12B shows the set-up time as a function of mesh resolution. Each barycentric refinement increases the set-up time by approximately one order of magnitude. Note: this increase is solely due to increased FEM and ADM run times as the rank required for a given accuracy does not increase.

After the set-up time of 8.5 – 10.5 hours, the reconstruction time to predict the E-field over the whole brain for a fixed coil placement is 0.57 – 0.85 seconds (using an Intel Core i7-10700 CPU, 2.90GHz) (meshed with ≈ 1.5 Million tetrahedra or ≈ 4.5 Million rows in the matrix ***A***.). In many applications, only the E-field on the cortex or on a small ROI is required. The reconstruction time varies linearly with the number of E-field samples. As such, to compute the E-field in a cortical surface consisting of ≈ 225*k* triangles takes ≈ 80 – 100 ms on the same CPU. Therefore, to compute the E-field on a coarse mesh consisting of ≈ 120,000 tetrahedra or ≈ 25,000 vertices, it would take approximately 17-22 ms or 55 - 60 ms, respectively, on the same CPU. These estimates are based on linear interpolation upon the current resolution of the mesh in this paper. As such, significant additional gains can be achieved by using lower-resolution meshes like in [21] & [22].

Table 3 provides CPU and GPU runtimes for the set-up and reconstruction of the E-field over the whole brain (consisting of 1.5 Million tetrahedra or, 4.5 Million matrix rows or, 325k vertices) and cortical surface (consisting of 240k triangular facets or, 720k matrix rows or, 120k vertices) for a range of CPUs, GPUs, and operating systems with a fixed rank, *r* = 110. For three different CPUs (a very low-end CPU-2 to a moderately high-end CPU-3), the average times are mentioned across five subjects for 1000 random coil placements. For the whole brain, the set-up time varies from 14.51-22.61 ms and the reconstruction time varies from 0.57-1.42 seconds for the E-field along 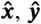, and 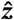 directions. The corresponding times for the cortical surface are mentioned in the Table. Furthermore, a mid-level Quadro RTX A6000 GPU takes 6.4 ms and 1.98 ms in the set-up and reconstruction stage, and a moderately new-generation GeForce RTX 3080 GPU takes only 1.21 ms and 0.33 ms, respectively. Irrespective of the hardware and software configuration, the reconstruction time involves the multiplication of a 110-dimensional vector with a number of E-field samples by a 110-dimensional dense matrix. These indicate that PMD could be used to determine the E-field in coil placement optimization applications rapidly.

**Table 3.** Average simulation time to predict the E-field for rank=110 in different hardware for a single coil placement across all tetrahedra inside the ROI; along x, y, and z direction, averaged across 5 subjects

| ROI | Simulation Stage | CPU-1<br>(Windows)<sup>1</sup> | CPU-2<br>(Centos)<sup>2</sup> | CPU-3<br>(Ubuntu)<sup>3*</sup> | GPU-1<sup>4</sup> | GPU-2<sup>5</sup> |
| --- | --- | --- | --- | --- | --- | --- |
| Brain | Set-up Time(ms)<sup>♦</sup> | 22.61 | 21.42 | 14.51 | — | — |
|  | Reconstruction Time(ms)<sup>♦♦</sup> | 1420.50 | 847 | 572 | 6.4 | 1.98 |
| Cortex | Set-up Time(ms)<sup>♦</sup> | 136.21 | 131.54 | 86.57 | — | — |
|  | Reconstruction Time(ms)<sup>♦♦</sup> | 287.87 | 143 | 99 | 1.21 | 0.33 |

<sup>1</sup> CPU-1 : Intel Xeon Gold 5220R CPU, 2.20GHz, dual processor, 24 Cores.
<sup>2</sup> CPU-2 : AMD Rome CPU, 2.0GHz, 128 Cores, Cluster CPU.
<sup>3</sup> CPU-3 : Intel Core i7-10700 CPU, 2.90GHz, 8 Cores.
<sup>4</sup> GPU-1 : NVIDIA Quadro RTX A6000, 48GB, 10752 Cuda Cores.
<sup>5</sup> GPU-2 : NVIDIA GeForce RTX 3080, 8GB, 8960 Cuda Cores.
\* Approximated by interpolation
♦ This is the total time for the algorithm to predict the E-field in a compressed form (as **Q** and **B**).
♦♦ This is the total time of reconstruction for the E-field from the compressed form, averaged across 1000 coil placements over each subject.

Fig. 13 shows the required peak memory for the algorithm as a function of rank. The required memory increases with PMD rank because additional E-field modes are required. Additionally, the memory required for the ‘QR decomposition’ in the algorithm increases (section 2.3). A single FEM simulation result (The E-field over the whole brain) consists of ≈ 1.5 Million tetrahedrons and hence, ≈ 4.5 Million matrix entries per mode. The memory requirement can be reduced further by storing each basis-mode and basis-coil separately in the second and fourth stages of the algorithm, respectively. Note: this memory also includes the overhead introduced by the MATLAB platform. Expanded CPU runtime and a summary of previously used fast E-field analysis platform results are provided in the supplementary file.

**Figure 13:**
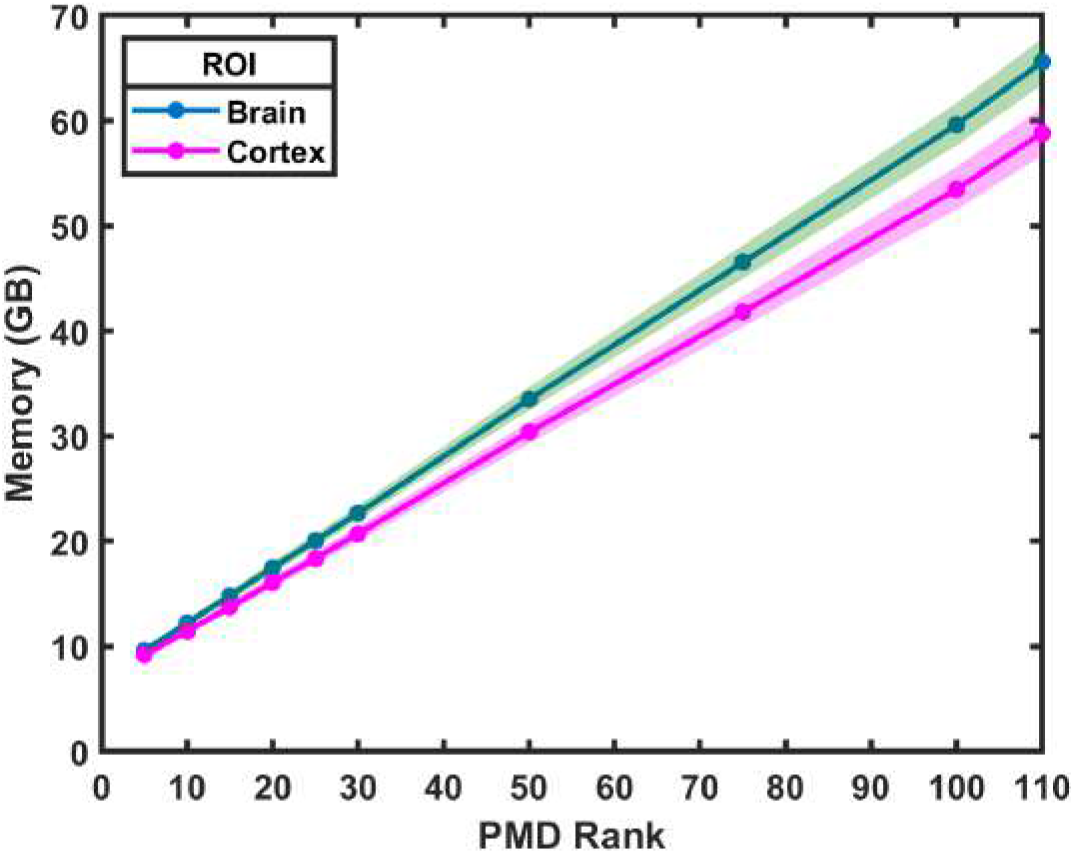
Required memory for the PMD algorithm when the ROI is either the whole brain or the cortex. The results are shown with a confidence interval, calculated across five subjects. The algorithm requires 60-65 GB peak memory for rank, *r* = 110 during run-time.

### 3.6. Application of the PMD for group level optimum coil placement

Here the average E-field in an ROI centered about MNI coordinate (−30,43,23) and having a diameter of 20 mm is considered. This location typically corresponds to the left Dorsolateral Prefrontal Cortex (Left-DLPFC). The average E-field in the ROI for coil placements on the EEG coordinate locations is shown in Fig. 14. (Note: for each coil placement we report the average E-field across all orientations.) The ‘F3’ electrode coordinate shows the highest Efield across individuals. However, the individual optimal varies as much as 37 mm from this location. Furthermore, the optimal coil placement across individuals is as much as 38 mm away from the ‘F3’ coordinate location. Detailed statistical distributions are shown in the supplementary Fig. S2.

**Figure 14:**
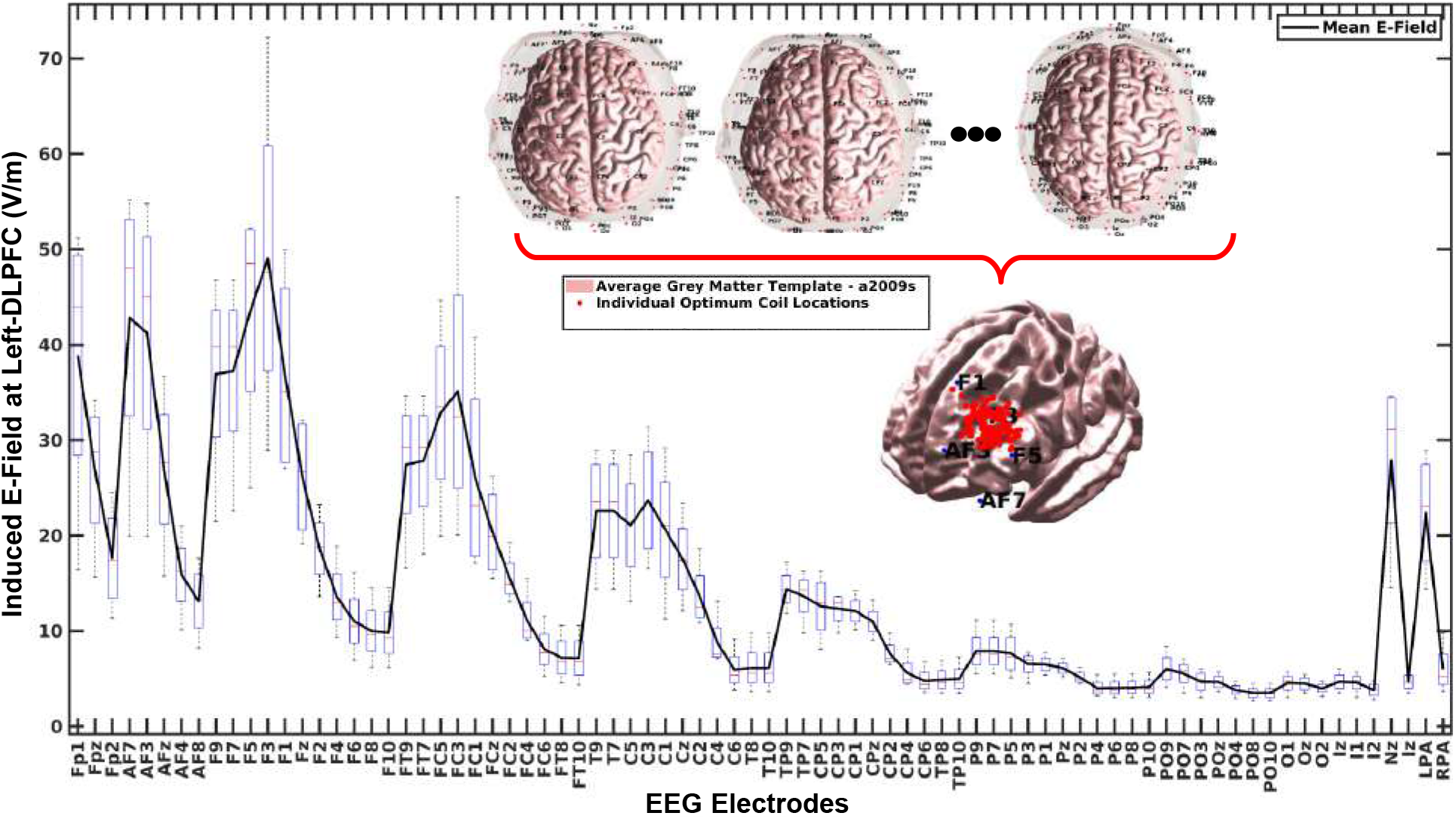
Average ROI E-field for different TMS coil placements. The distribution of the E-field across 200 subjects shows the highest E-field is expected for the ‘F3’ coil placement. Individual optimum coil placements are all distinct and group-level optimal coil placement is as much as 55 mm away from the ROI and at most 28 mm from the ‘F3’ coordinate.

## 4. Discussion

The PMD approximation technique enables the rapid prediction of induced E-field in the whole brain with a large number of coil placements over the scalp with an initial set-up time of only ≈ 9.5 hours. The validation results indicate that a rank of *r* = 110 is sufficient for accurately predicting the induced E-field for up to 1 Million coil placements. For example, the observed mean value of the ***RMSRE*** across individuals is ≈ 1%. Moreover, the proposed ***PMDRES***_2_ and ***PMDRES**_F_* metrics provide an adequate estimate for the actual ***RMSRE*** error. All of these results were validated in 200 subjects.

A detailed study on the proposed methodology shows that the required PMD rank is insensitive to the density of coil placements and head model mesh. This is because the green’s function linking sources on a scalp surface to the E-field observable is compressible, and the rank only depends on the size of the source domain and its separation from the observable [24]. As such the total rank is only dependent on the error required and the size of the region covered by all coil placements. We believe that these boundary coil location errors are due to a truncation effect. Specifically, the dipole models of coils oftentimes overlap, thereby, resulting in a prioritizing of the accurate representation of coil placements that result in the most overlap. The dipoles on the boundaries of the coil placement mesh do not have as many overlapping dipoles as those interiors. We anticipate that the increased error in the boundaries can be reduced through preconditioning, in other words, increasing the relative weighting of the matrix entries corresponding to boundary coil positions in the matrix.

As the mesh density is increased, the required rank to reach 2 % ***PMDRES***_2_ error decreased marginally. For example, to reach 2 % error we need a rank of 110, however, on the twice barycentrically refined mesh we only require a rank of 100. This is likely due to the accuracy of the simulation increasing. The matrix that we compressed is an approximation to Laplace’s equation. As such, its singular values and vectors consist of Laplace operator parts and numerical artifacts. We believe that the numerical artifacts result in slowed convergence of the PMD. As such, when we increased the fidelity of the results, the rank required for a given accuracy decreased. The PMD algorithm is agnostic to E-field solver, a higher order FEM or BEM could be used to improve convergence without the need for barycentric refinement [12].

A comprehensive study of the speed of the method shows that the reconstruction time required is only ≈ 0.57 seconds per coil placement in a very low-end CPU. The necessary computational resources for this process are also minimal, requiring at most a standard laptop with standard configurations. An exhaustive grid search over all possible coil placement configurations with the same computational resources would have taken ≈ 5 years for a single subject. This platform can be efficiently used to find group-level optimum coil placements. Furthermore, the reconstruction stage can be executed in a GPU enabling the computation of the E-field distributions in under 2-3 milliseconds enabling the use of this method to determine optimum coil placement on the fly for rapid coil placement reconfiguration.

## 5. Conclusion

We presented a method for compressing a matrix that encodes the relationship between coil position to E-field. Using this method the E-field in the brain can be determined in 0.57 seconds and 0.33 milliseconds using a CPU or a GPU, respectively, over the whole brain. The PMD compression takes about 9.5 hours of pre-processing time (set-up time). This framework enables the practical E-field-informed group-level coil placement optimization. Future work will include defining projectors that enable the use of PMD compressed matrices for real-time E-field dosimetry of TMS for absolutely arbitrary coil placement.

## Credit authorship contribution statement

### Nahian Ibn hasan

Conceptualization, Methodology, Software, Validation, Formal analysis, Investigation, Data curation, Writing - original draft, Writing - review & editing, Visualization.

### Dezhi Wang

Conceptualization, Methodology, Software, Investigation.

### Luis J. Gomez

Funding acquisition, Conceptualization, Supervision, Methodology, Software, Validation, Formal analysis, Investigation, Writing - review & editing, Funding acquisition.

## Data and Code Availability Statement

Data and Code Availability Statement: In-house codes are openly available at this link - **Link will be available after publication**

## Supplementary Material

Supplementary materials can be found in the online version of this research article.

## Supplementary Material

### S1. Similarity Metrics

Fig. S1(A-D) shows different similarity metrics (Pearson Correlation Coefficient (PCC), Cosine Similarity(CS), Triangle Area Similarity(TS), and Sector Area Similarity(SS), respectively) between the ground truth and approximated E-field for PMD rank *r* =110 across 1000 random coil placements according to equations S1e, S1f, S1g and S1h. Both PCC and CS achieve a near-perfect similarity (close to 1) between actual and predicted E-fields. PCC measures the similarity in terms of the magnitude of the E-fields whereas CS measures the directional similarity in between them without considering the magnitude. On the other hand, TS measures the similarity between actual and predicted E-fields based on their directions, magnitudes, and Euclidean error (ED) in between them [40]. A very small value of the TS metric means the two quantities are very similar. In our case, this value is of the order of 10^−7^. However, TS does not consider the difference in magnitude of the two quantities. Therefore, we use another metric, SS [40], which also considers the magnitude error (MD) along with direction, individual magnitudes, and ED. An SS score of 0 means both ED and MD are zero and there is a perfect match between ground truth and predicted E-field. In our case, this score is of the order of 10^−17^, a near-perfect score.

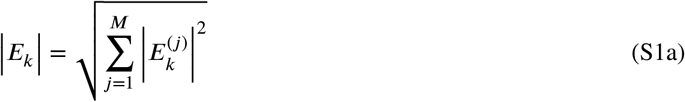

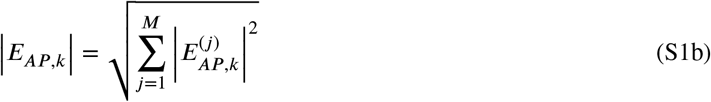

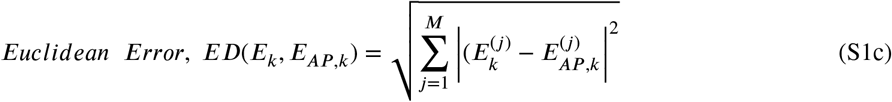

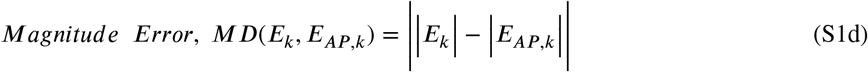

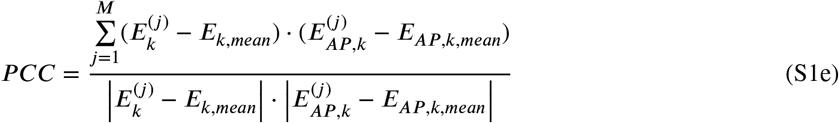

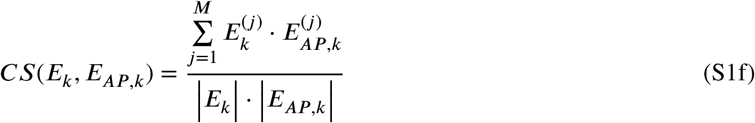

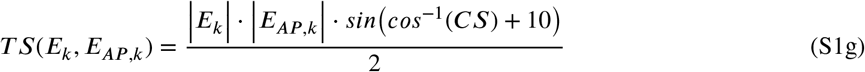

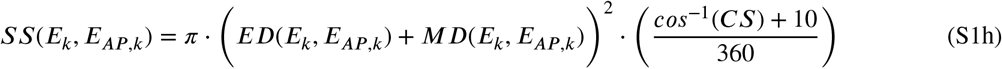

**Figure S1:**
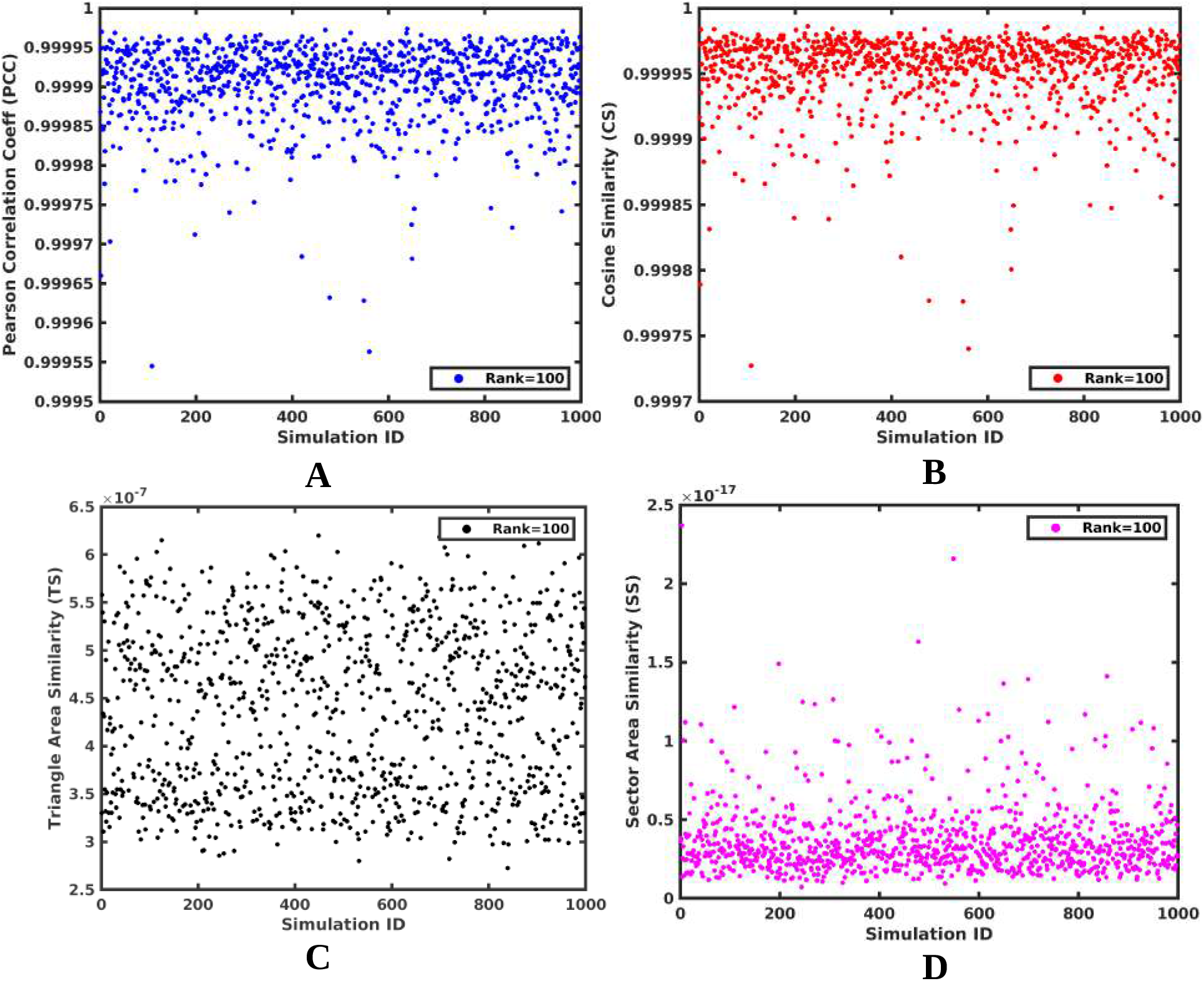
Different similarity metrics between the ground truth and approximated E-field for rank 100 across 1000 random coil placements.

### S2. E-Field analysis platforms

**Table S1.**
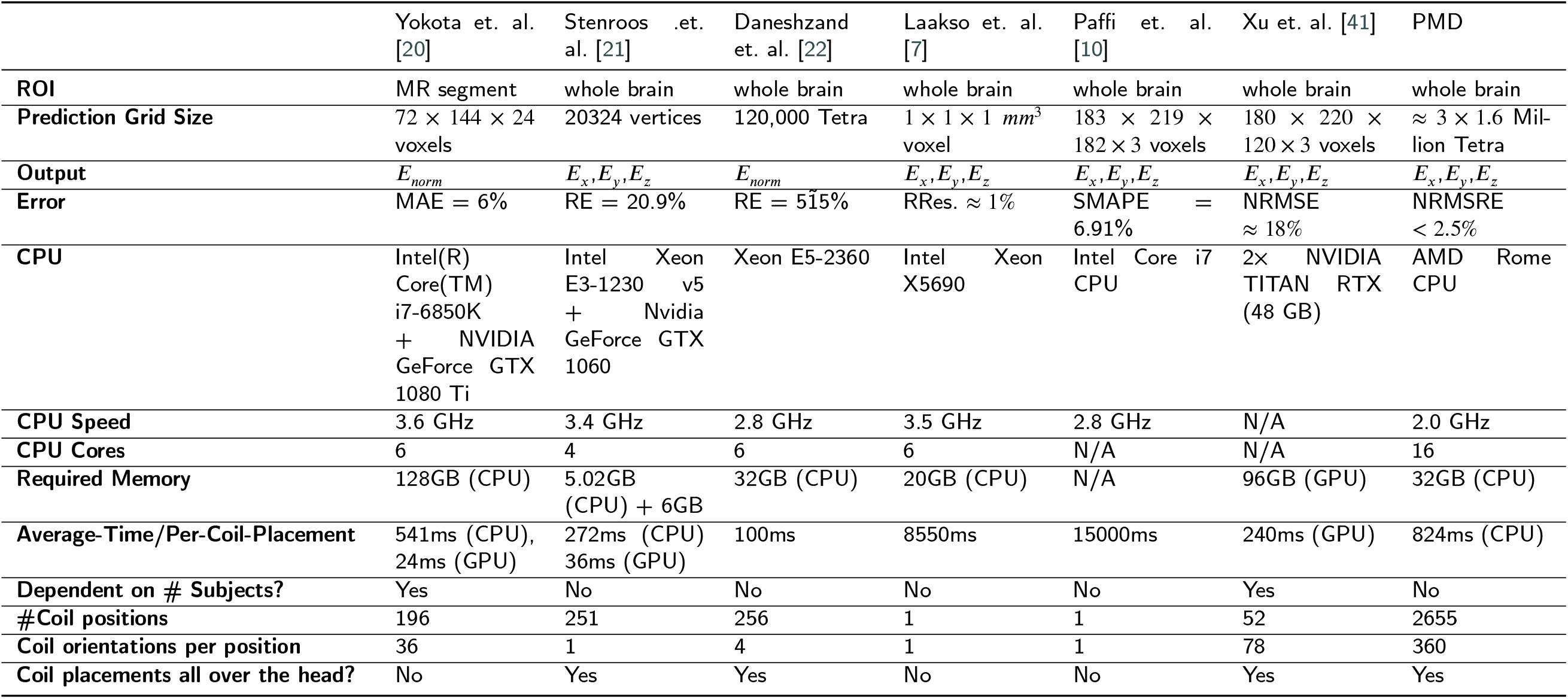
Simulation time and error comparison across different methods

### S3. Statistical distribution of individual, group-level and F3-protocol based optimum coil placements

**Figure S2:**
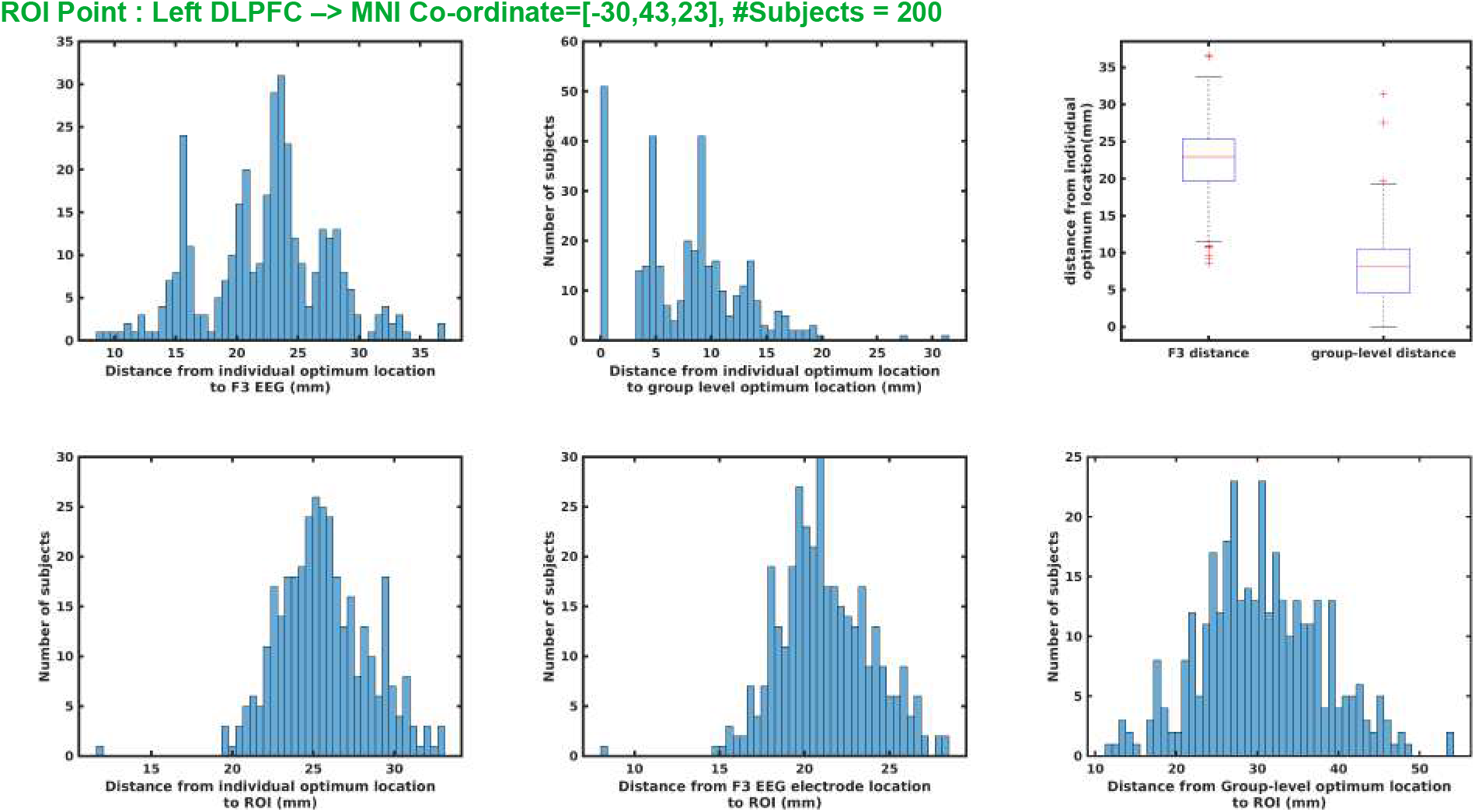
Statistical distribution of individual, group-level, and F3-protocol-based optimum coil placements and the distance among them across 200 subjects in the ROI of Left-DLPFC.

