## supplementary PDF for "Fast And Accurate Population Level Transcranial Magnetic Stimulation via Low-Rank Probabilistic Matrix Decomposition (PMD)"

### Supplementary Material

#### S1. Similarity Metrics

Fig. S1(A-D) shows different similarity metrics (Pearson Correlation Coefficient (PCC), Cosine Similarity(CS), Triangle Area Similarity(TS), and Sector Area Similarity(SS), respectively) between the ground truth and approximated E-field for PMD rank $r = 110$ across 1000 random coil placements according to equations S1e, S1f, S1g and S1h. Both PCC and CS achieve a near-perfect similarity (close to 1) between actual and predicted E-fields. PCC measures the similarity in terms of the magnitude of the E-fields whereas CS measures the directional similarity in between them without considering the magnitude. On the other hand, TS measures the similarity between actual and predicted E-fields based on their directions, magnitudes, and Euclidean error (ED) in between them [40]. A very small value of the TS metric means the two quantities are very similar. In our case, this value is of the order of $10^{-7}$. However, TS does not consider the difference in magnitude of the two quantities. Therefore, we use another metric, SS [40], which also considers the magnitude error (MD) along with direction, individual magnitudes, and ED. An SS score of 0 means both ED and MD are zero and there is a perfect match between ground truth and predicted E-field. In our case, this score is of the order of $10^{-17}$, a near-perfect score.

$$\left|E_k\right| = \sqrt{\sum_{j=1}^{M} \left|E_k^{(j)}\right|^2} \tag{S1a}$$

$$\left|E_{AP,k}\right| = \sqrt{\sum_{j=1}^{M} \left|E_{AP,k}^{(j)}\right|^2} \tag{S1b}$$

$$Euclidean\ \ Error,\ ED(E_k, E_{AP,k}) = \sqrt{\sum_{j=1}^{M} \left|(E_k^{(j)} - E_{AP,k}^{(j)}\right|^2} \tag{S1c}$$

$$Magnitude\ \ Error,\ MD(E_k, E_{AP,k}) = \left|\left|E_k\right| - \left|E_{AP,k}\right|\right| \tag{S1d}$$

$$PCC = \frac{\sum_{j=1}^{M} (E_k^{(j)} - E_{k,mean}) \cdot (E_{AP,k}^{(j)} - E_{AP,k,mean})}{\left|E_k^{(j)} - E_{k,mean}\right| \cdot \left|E_{AP,k}^{(j)} - E_{AP,k,mean}\right|} \tag{S1e}$$

$$CS(E_k, E_{AP,k}) = \frac{\sum_{j=1}^{M} E_k^{(j)} \cdot E_{AP,k}^{(j)}}{\left|E_k\right| \cdot \left|E_{AP,k}\right|} \tag{S1f}$$

$$TS(E_k, E_{AP,k}) = \frac{\left|E_k\right| \cdot \left|E_{AP,k}\right| \cdot sin\left(cos^{-1}(CS) + 10\right)}{2} \tag{S1g}$$

$$SS(E_k, E_{AP,k}) = \pi \cdot \left(ED(E_k, E_{AP,k}) + MD(E_k, E_{AP,k})\right)^2 \cdot \left(\frac{cos^{-1}(CS) + 10}{360}\right) \tag{S1h}$$

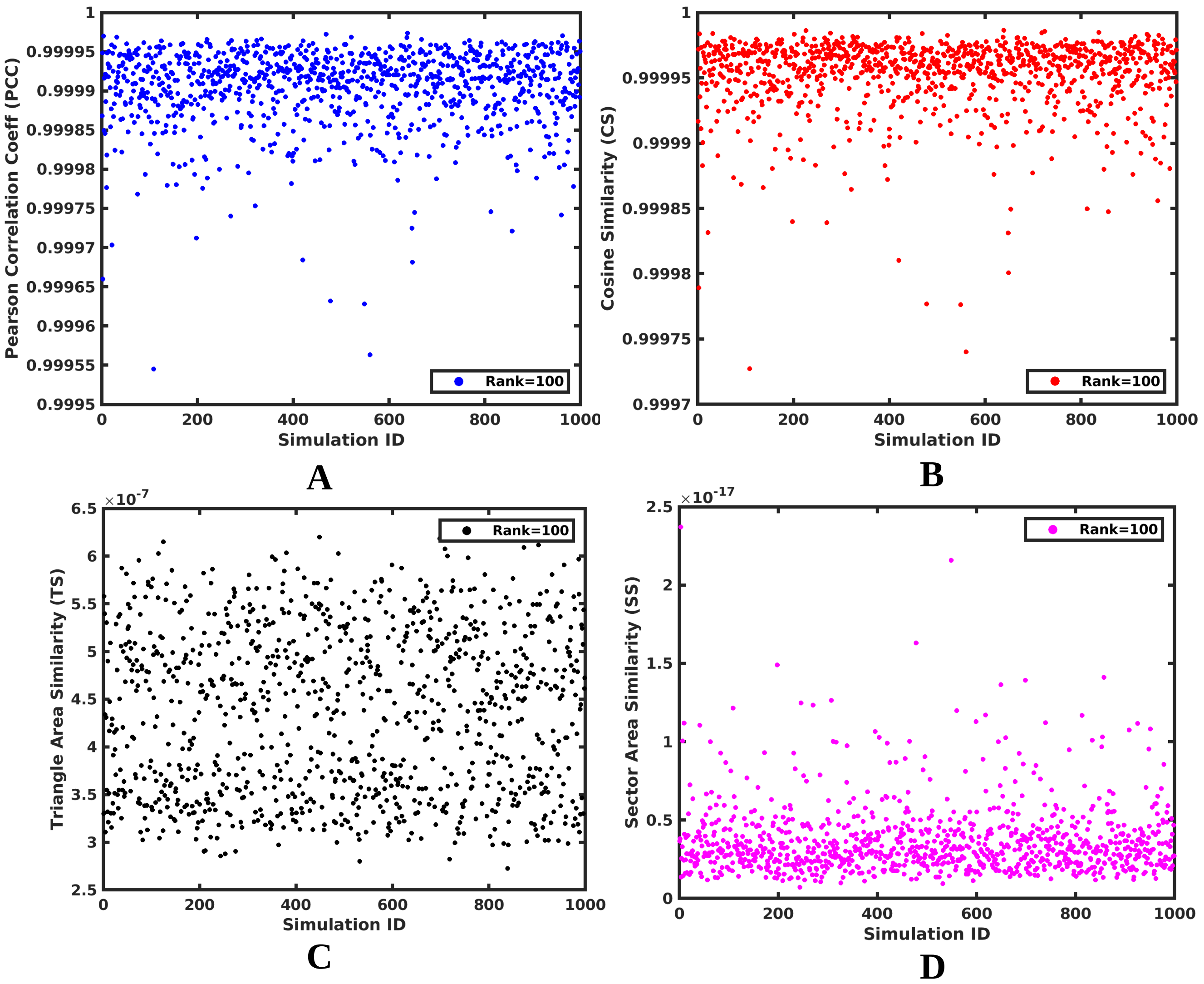


**Figure S1:** Different similarity metrics between the ground truth and approximated E-field for rank 100 across 1000 random coil placements.

#### S2. E-Field analysis platforms

**Table S1**
Simulation time and error comparison across different methods

| | Yokota et. al. [20] | Stenroos .et. al. [21] | Daneshzand et. al. [22] | Laakso et. al. [7] | Paffi et. al. [10] | Xu et. al. [41] | PMD |
|---|---|---|---|---|---|---|---|
| **ROI** | MR segment | whole brain | whole brain | whole brain | whole brain | whole brain | whole brain |
| **Prediction Grid Size** | $72 \times 144 \times 24$ voxels | 20324 vertices | 120,000 Tetra | $1 \times 1 \times 1$ $mm^3$ voxel | $183 \times 219 \times 182 \times 3$ voxels | $180 \times 220 \times 120 \times 3$ voxels | $\approx 3 \times 1.6$ Million Tetra |
| **Output** | $E_{norm}$ | $E_x, E_y, E_z$ | $E_{norm}$ | $E_x, E_y, E_z$ | $E_x, E_y, E_z$ | $E_x, E_y, E_z$ | $E_x, E_y, E_z$ |
| **Error** | MAE = 6% | RE = 20.9% | RE = 5$\tilde{1}$5% | RRes. $\approx 1\%$ | SMAPE = 6.91% | NRMSE $\approx 18\%$ | NRMSRE $< 2.5\%$ |
| **CPU** | Intel(R) Core(TM) i7-6850K + NVIDIA GeForce GTX 1080 Ti | Intel Xeon E3-1230 v5 + Nvidia GeForce GTX 1060 | Xeon E5-2360 | Intel Xeon X5690 | Intel Core i7 CPU | 2× NVIDIA TITAN RTX (48 GB) | AMD Rome CPU |
| **CPU Speed** | 3.6 GHz | 3.4 GHz | 2.8 GHz | 3.5 GHz | 2.8 GHz | N/A | 2.0 GHz |
| **CPU Cores** | 6 | 4 | 6 | 6 | N/A | N/A | 16 |
| **Required Memory** | 128GB (CPU) | 5.02GB (CPU) + 6GB | 32GB (CPU) | 20GB (CPU) | N/A | 96GB (GPU) | 32GB (CPU) |
| **Average-Time/Per-Coil-Placement** | 541ms (CPU), 24ms (GPU) | 272ms (CPU) 36ms (GPU) | 100ms | 8550ms | 15000ms | 240ms (GPU) | 824ms (CPU) |
| **Dependent on # Subjects?** | Yes | No | No | No | No | Yes | No |
| **#Coil positions** | 196 | 251 | 256 | 1 | 1 | 52 | 2655 |
| **Coil orientations per position** | 36 | 1 | 4 | 1 | 1 | 78 | 360 |
| **Coil placements all over the head?** | No | Yes | Yes | No | No | Yes | Yes |

#### S3. Statistical distribution of individual, group-level and F3-protocol based optimum coil placements

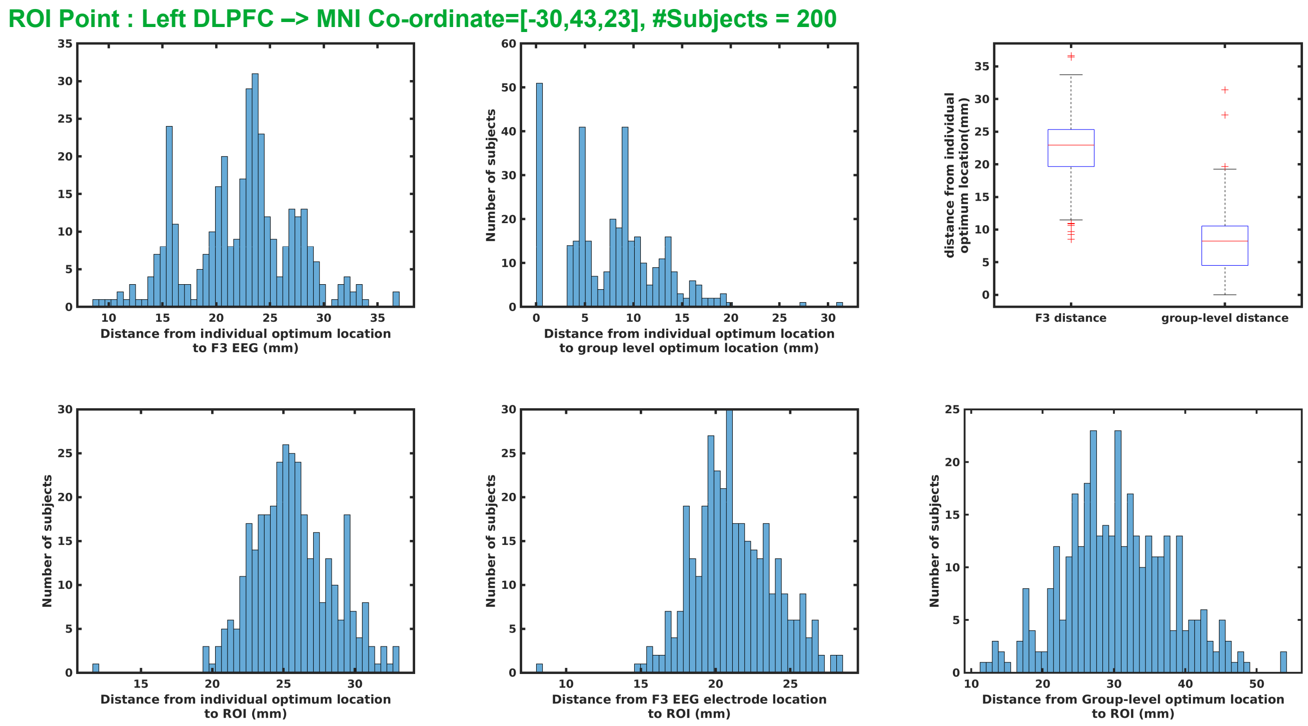


**Figure S2:** Statistical distribution of individual, group-level, and F3-protocol-based optimum coil placements and the distance among them across 200 subjects in the ROI of Left-DLPFC.
